# Pharmacological chaperones restore proteostasis of epilepsy-associated GABA_A_ receptor variants

**DOI:** 10.1101/2023.04.18.537383

**Authors:** Ya-Juan Wang, Hailey Seibert, Lucie Y. Ahn, Ashleigh E. Schaffer, Ting-Wei Mu

## Abstract

Recent advances in genetic diagnosis identified variants in genes encoding GABA_A_ receptors as causative for genetic epilepsy. Here, we selected eight disease-associated variants in the α1 subunit of GABA_A_ receptors causing mild to severe clinical phenotypes and showed that they are loss of function, mainly by reducing the folding and surface trafficking of the α1 protein. Furthermore, we sought client protein-specific pharmacological chaperones to restore the function of pathogenic receptors. Applications of positive allosteric modulators, including Hispidulin and TP003, increase the functional surface expression of the α1 variants. Mechanism of action study demonstrated that they enhance the folding and assembly and reduce the degradation of GABA_A_ variants without activating the unfolded protein response in HEK293T cells and human iPSC-derived neurons. Since these compounds cross the blood-brain barrier, such a pharmacological chaperoning strategy holds great promise to treat genetic epilepsy in a GABA_A_ receptor-specific manner.

## Introduction

Gamma-aminobutyric acid type A (GABA_A_) receptors are the primary inhibitory neurotransmitter-gated ion channels in the mammalian central nervous system ^1^. They mediate the majority of the postsynaptic inhibitory action to maintain the excitation-inhibition balance under physiological conditions. Recent advance in genetics has identified an increasing number of epilepsy-associated genes, among which, genes encoding GABA_A_ receptors are prominent causative factors ^2–4^. GABA_A_ receptors are pentamers, which assemble from specific combinations of eight subunit classes: α1-6, β1-3, γ1-3, δ, ε, θ, π, and ρ1-3. GABA_A_ receptors are distributed throughout the brain regions, with the most common subtype containing two α1 subunits, two β2 subunits, and one γ2 subunit ^5^. Currently, hundreds of clinical variants have been identified in GABA_A_ receptor subunits (www.clinvar.com), including the major α1, β2, β3, and γ2 subunits ^6, 7^. The majority of GABA_A_ receptor variants cause their loss of function, although recent studies support some gain-of-function variants ^8^. Despite the development of numerous anti-seizure drugs, about one third of epilepsy patients are resistant to current drug treatment ^9^, and many of them have a genetic diagnosis ^10^. Therefore, there is an urgent need to develop a new therapeutic strategy to treat genetic epilepsy resulting from functional defects of GABA_A_ receptors.

Proteostasis maintenance is critical for the function of GABA_A_ receptors ^11^. Individual subunit of GABA_A_ receptors needs to fold into its three-dimensional native structure and assembles with other subunits to form a heteropentamer in the endoplasmic reticulum (ER). Only properly assembled pentameric receptors exit the ER and traffic through the Golgi and onward to the plasma membrane to perform their function. Misfolded or unassembled subunits are retained in the ER; they undergo additional folding/assembly cycles or are targeted to cellular degradation pathways, including ER-associated degradation (ERAD) ^12^. The proteostasis network orchestrates the folding, assembly, trafficking, and degradation of GABA_A_ receptors. Recently, our quantitative proteomics identified a proteostasis network for GABA_A_ receptors, which enabled our efforts to further elucidate their biogenesis pathways ^13^. Upon the synthesis of GABA_A_ receptor subunits, the ER membrane protein complex (EMC), including EMC3 and EMC6, facilitates the insertion of the transmembrane domain of GABA_A_ receptors into the ER membrane ^14^. Then BiP (binding immunoglobulin protein) and calnexin in the ER act as pro-folding chaperones to promote the productive folding of GABA_A_ receptors ^15, 16^. Furthermore, Hsp47 (heat shock protein 47) acts after BiP to interact with relatively folded subunits to enhance the subunit assembly process into pentameric receptors in the ER ^17^. Then the receptors engage the trafficking receptors, such as LMAN1 (lectin, mannose binding 1) ^18^, for transport to the Golgi and plasma membrane.

One major disease-causing mechanism for the loss of function of GABA_A_ receptors is that pathogenic variants lead to receptor trafficking deficiencies due to defective protein conformations ^6, 19^. Such epilepsy-associated GABA_A_ receptor subunit variants cause protein misfolding and/or disrupt the subunit-subunit assembly process in the ER, resulting in a decrease in the surface expression of the pentameric receptors. A promising therapeutic strategy to restore the function of misfolding-prone variants is to correct their folding, trafficking, and thereby function. This strategy has dual roles: first, the surface presence of GABA_A_ receptors will be restored; second, the restored GABA_A_ receptors on the plasma membrane will be accessible by GABAergic signaling molecules that act on the plasma membrane. Additionally, this strategy is not limited to specific GABA_A_ variants, since it targets the overall folding and trafficking of GABA_A_ receptors; furthermore, this strategy can achieve selectivity for destabilized variants over wild type proteins. Two mechanistically distinct classes of small molecules that are used to correct protein conformational defects are called proteostasis regulators and pharmacological chaperones ^20, 21^. Proteostasis regulators promote the proteostasis network capacity by transcriptional and translational re-programming of the network components ^22, 23^. Recently, we have identified a number of proteostasis regulators with different mechanisms of action that can correct the function of several GABA_A_ variants ^15, 16, 24–26^. For example, pharmacologically activating the ATF6 (Activating transcription factor 6) branch of the unfolded protein response selectively corrects the folding, trafficking, and thus function of destabilized GABA_A_ receptor variants ^26^. As distinct from proteostasis regulators, pharmacological chaperones directly bind their target proteins to stabilize them; the enhanced stability will promote the cellular trafficking of the target proteins to their functional location. The advantage of pharmacological chaperones is that they can be specific to their target proteins. The pharmacological chaperoning strategy has been applied to restore the trafficking of defective lysosomal storage enzymes ^27^ and certain G-protein-coupled receptors (GPCRs) ^28^. However, their application on trafficking deficient GABA_A_ receptor variants has not been reported.

We seek GABA_A_ receptor-specific pharmacological chaperones. The pharmacology of GABA_A_ receptors has been studied extensively ^29, 30^. Numerous agonists, antagonists, and allosteric modulators, such as benzodiazepines, have been identified. Since they directly bind GABA_A_ receptors, they have the promise to act as pharmacological chaperones to promote the trafficking of GABA_A_ variants. The structures of pentameric GABA_A_ receptors have been well-characterized ^31, 32^. Each subunit has a long extracellular (or the ER lumenal) N-terminal domain (NTD), a transmembrane domain containing four transmembrane (TM) helices TM1-4, a short cytosolic loop between TM1 and TM2, a short extracellular (or the ER lumenal) loop between TM2 and TM3, a large cytosolic loop between TM3 and TM4, and a short extracellular (or the ER lumenal) C-terminus (**Supplementary Fig. S1a**). Each pentameric α1β2γ2 GABA_A_ receptor has two GABA binding sites in the NTD at the β2/α1 interfaces (**Supplementary Fig. S1b**): residues from the β2 subunit form the principal “positive” side, whereas residues from the α1 subunit form the complementary “negative” side of the ligand-binding pocket. In addition, each pentameric α1β2γ2 receptor has one classic high-affinity benzodiazepine binding site in the NTD at the α1+/γ2− interface (**Supplementary Fig. S1b**); recent cryogenic electron microscopy (cryo-EM) studies identified additional low-affinity benzodiazepine binding sites in the transmembrane domains at the subunit interfaces ^33, 34^.

In this study, we aimed to identify potent pharmacological chaperones to correct the function of epilepsy-associated GABA_A_ receptors. Our screen of known GABA_A_ receptor modulators discovered that hispidulin and TP003, which act on the benzodiazepine binding site in GABA_A_ receptors, effectively restored the function of several pathogenic variants causative for mild and severe clinical phenotypes. Both chemicals promoted surface trafficking of the GABA_A_ variants by enhancing subunit folding and assembly as well as reducing ERAD.

## Results

### The majority of selected disease-associated variants (DAVs) of the GABA_A_ receptor α1 subunits lead to reduced surface α1 protein levels

In this study, we focus on the α1 subunit of GABA_A_ receptors: a growing number of clinical variants (> 200 variants according to www.clinvar.com, including missense, frameshift, and nonsense variants) have been identified in this subunit to be associated with genetic epilepsy covering a broad phenotypic spectrum ^6, 7, 35, 36^. These variations are distributed throughout the protein sequence in the NTD, the transmembrane domain (TM1, TM2, TM3, and TM4), and the loops connecting TM helices (**Fig. 1a** and **Supplementary Fig. S2**). We used Rhapsody, which integrates both sequence evolution and structure- and dynamics-based data ^37^, to carry out *in silico* saturation mutagenesis analysis of the α1 subunit. According to the calculated pathogenic probability, most TM1, TM2, and TM3 positions are susceptible to disease-causing variations, whereas TM4 is less susceptible, possibly because the insertion of TM4 into the ER membrane occurs late in the biogenesis step. In addition, the ER lumenal loop between TM2 and TM3 and the NTD loop between β8 and β9 strands that is in proximity to the pre-TM1 region are susceptible to disease-causing variations (**Supplementary Fig. S2**). The currently known α1 missense variants were plotted using their predicted pathogenic probability against their primary protein sequence (**Fig. 1a** and **Supplementary Table S1**): 27 variants are predicted to be mild with a pathogenic probability of below 0.30, 61 variants are predicted to be moderate with a pathogenic probability of between 0.30 and 0.60, and 61 variants are predicted to be severe with a pathogenic probability of above 0.60. The spatial distribution of available positions of such missense variants was illustrated in the α1 subunit cryo-EM structure ^38^ (**Fig. 1b**).

**Fig 1.**
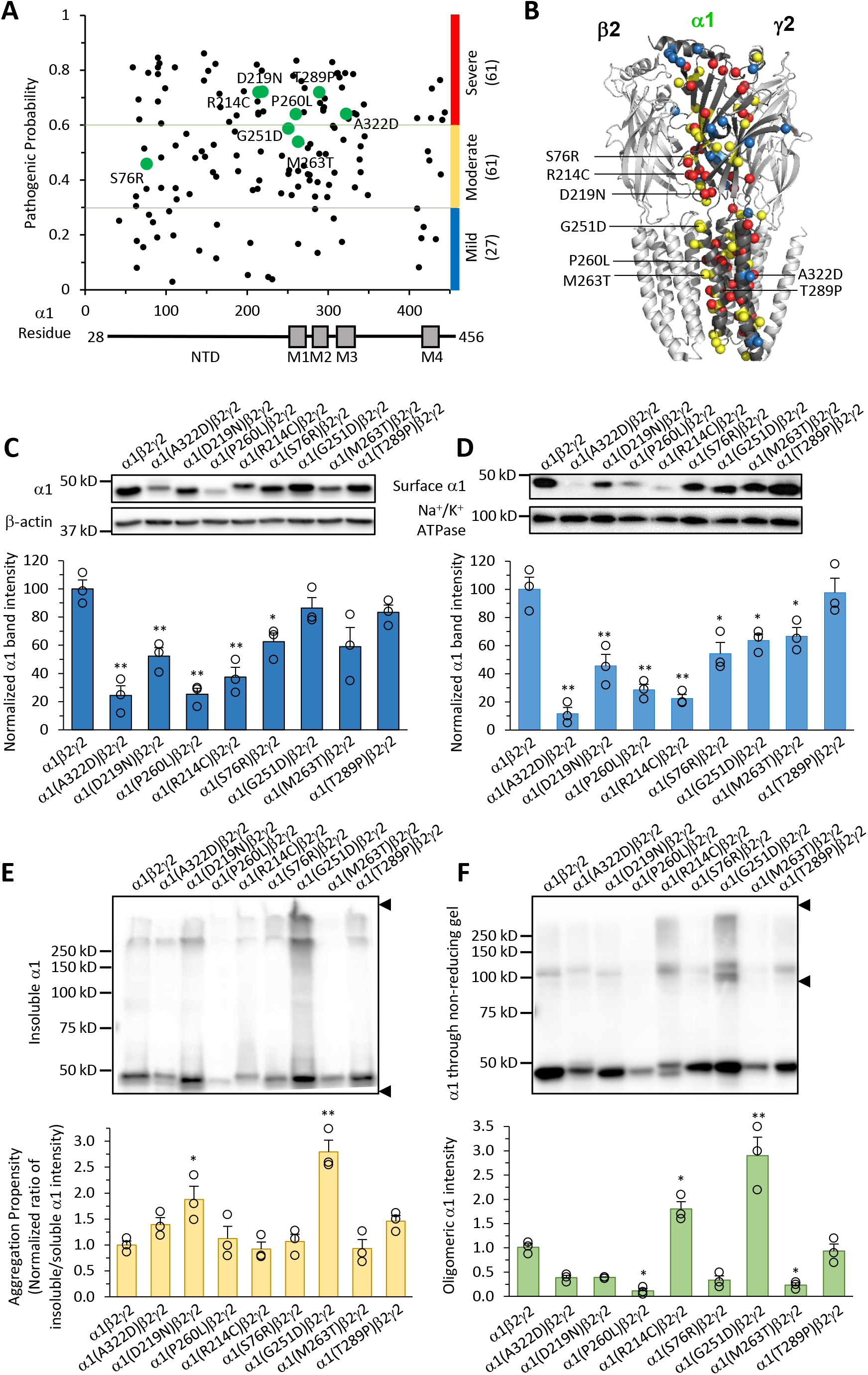
Molecular characterization of selected disease-associated variants (DAVs) of GABA_A_ receptor α1 subunits. **a** Pathogenic probability of 149 clinical variants of the GABA_A_ receptor α1 subunit is predicted using Rhapsody and plotted against its primary protein sequence. A schematic of the mature α1 sequence (residue 28-456) showing the N-terminal domain (NTD) and the transmembrane domain (M1-M4) is displayed on the bottom. The variants are classified into three categories: mild (pathogenic probability < 0.30), moderate (0.30 ≤ pathogenic probability < 0.60), and severe (pathogenic probability ≥ 0.60). Eight variants are selected for further characterization and colored in green. **b** The spatial distribution of the positions of α1 variants is illustrated in cryo-EM structure of α1β2γ2 GABA_A_ receptors (6X3S.pdb), rendered using PyMOL. The C_β_ of the residues (C_α_ in the case of glycine) are shown as spheres. Mild variants are colored in blue, moderate variants in yellow, and severe variants in red. Eight selected variants are labelled. **c** HEK293T cells were transfected with α1 (wild type or the indicated variants), β2, and γ2 at a ratio of 1:1:1. Forty-eight hours post transfection, cells were lysed with a lysis buffer containing 2 mM DDM. The total proteins were subjected to SDS-PAGE and Western blot analysis. β-actin serves as a total protein loading control. Quantification of the normalized α1 band intensity is shown on the bottom (n = 3). **d** Surface biotinylation assay was used to quantify the surface α1 protein level. Na^+^/K^+^-ATPase serves as a plasma membrane protein loading control. Quantification of the normalized surface α1 band intensity is shown on the bottom (n = 3). **e** Insoluble α1 fraction was generated from removing soluble α1 fraction as shown in **c** by extracting proteins with a lysis buffer containing 2 mM DDM; residual insoluble α1 was re-suspended with Laemmli sample buffer containing 2% SDS and subjected to Western blot analysis. The band intensity was quantified by including the entire lane as indicated within the two arrows. Quantification of the ratio of insoluble over soluble α1 as a measure of aggregation propensity is shown on the bottom (n = 3). **f** Non-reducing protein gel was used to determine the oligomerization of α1 subunits. Total proteins were extracted and resolved through non-reducing SDS-PAGE in the absence of a reducing reagent. Quantification of oligomeric α1, as indicated within the two arrows, is shown on the bottom (n = 3). Each data point is reported as mean ± SEM. One-way ANOVA followed by post-hoc Tukey test was used for statistical analysis. *, *p* < 0.05; **, *p* < 0.01. Also see **Supplementary Table S1** and **Supplementary Fig. S2**.

We selected eight DAVs in the α1 subunit to evaluate their proteostasis deficiency, namely S76R, R214C, D219N, G251D, P260L, M263T, T289P, and A322D, by considering the following two criteria. First, these variants, possessing different predicted pathogenic probability (**Fig. 1a**), cover a broad phenotypic spectrum: S76R ^39^, R214C ^40, 41^, G251D ^39^, P260L ^42^, M263T ^42^, and T289P ^39^ cause severe epileptic encephalopathies, such as Dravet and Dravet-like syndrome and West syndrome; D219N ^43^ and A322D ^44^ cause milder idiopathic generalized epilepsy (IGE), such as juvenile myoclonic epilepsy (JME). Second, these variants represent diverse spatial distribution in the α1 subunit: S76R, R214C, and D219N are located in the NTD, G251D, P260L, and M263T are in TM1, T289P is in TM2, and A322D is in TM3 (**Fig. 1b** and **Supplementary Fig. S2**).

We investigated the effect of the eight DAVs on the total protein and surface protein levels of the α1 subunits. We used HEK293T cells to exogenously express α1 DAVs with wild type β2 and γ2 subunits since HEK293T cells do not possess endogenous GABA_A_ receptors. Western blot analysis demonstrated that five variations (S76R, R214C, D219N, P260L, and A322D) reduced the total α1 subunit protein levels significantly compared to wild type, suggesting an excessive degradation of such DAVs (**Fig. 1c**). Furthermore, surface biotinylation assay showed that seven variations (S76R, R214C, D219N, G251D, P260L, M263T, and A322D) reduced the surface α1 protein levels significantly (**Fig. 1d**), consistent with previous reports that R214C ^40^, D219N ^43^, and A322D ^45^ compromised the trafficking of α1 DAVs to the plasma membrane. The most dramatic decrease was observed in R214C, P260L, and A322D variations, as 37.5%, 25.2%, and 24.3% of total wild type α1 (**Fig. 1c**), and 22.4%, 28.7%, and 11.7% of surface wild type α1 (**Fig. 1d**), respectively.

Furthermore, we evaluated the aggregation propensity of the α1 DAVs by using detergent solubility assay (**Fig. 1e**). After the solubilization of the cell lysates in n-dodecyl-β-maltoside (DDM), a non-ionic detergent that can effectively solubilize GABA_A_ receptors, the pellet fraction was re-suspended with 2% SDS and visualized through SDS-PAGE and Western blot analysis. The ratio of the pellet fraction / supernatant fraction was used as a measure of the aggregation propensity. Intriguingly, the G251D variation led to the most dramatic increase of the aggregation propensity by 2.8-fold compared to wild type α1 (**Fig. 1e**, cf. lane 7 to 1). In addition, we used non-reducing protein electrophoresis, which preserves GABA_A_ receptor subunit-subunit interactions due to the absence of reducing agents, to evaluate the oligomerization states of α1 DAVs (**Fig. 1f**). Notably, the G251D variation accumulated substantially more high molecular weight α1 oligomers compared to wild type α1 (**Fig. 1f**, cf. lane 7 to 1); additionally, the R214C variation resulted in more α1 oligomers (**Fig. 1f**, cf. lane 5 to 1), possibly due to the introduction of an extra cysteine for potential crosslinking reactions.

Our data demonstrated that reduced surface trafficking was observed for seven among eight selected α1 DAVs, indicating that trafficking deficiency is one major disease-causing mechanism for GABA_A_ receptor variants. In addition, the G251D variation aggravated its cellular oligomerization and aggregation, whereas S76R, R214C, D219N, P260L, and A322D led to their excessive cellular degradation.

### All selected **α**1 DAVs reveal loss-of-function effects as demonstrated by whole-cell patch-clamp recordings

Presumably, pathogenic GABA_A_ receptor variants cause their loss of function and thus reduced inhibitory neuronal activity and epilepsy syndromes. However, recent studies reported gain-of-function variants in β3 subunits of GABA_A_ receptors ^8^. Therefore, we further investigated the effect of these α1 DAVs on the functionality of the GABA_A_ receptors using whole-cell patch-clamp recordings in HEK293T cells. GABA (100 μM)-induced peak current amplitudes were significantly reduced for all eight α1 DAVs compared to wild type receptors (**Fig. 2a**), clarifying loss-of-function effects for these variants. Wild type receptors exhibited a peak current amplitude of 6.3 ± 0.4 nA, and the most dramatic current decreases were observed in the variants of A322D (1.6 ± 0.2 nA) and P260L (1.9 ± 0.3 nA) (**Fig. 2a**).

**Fig 2.**
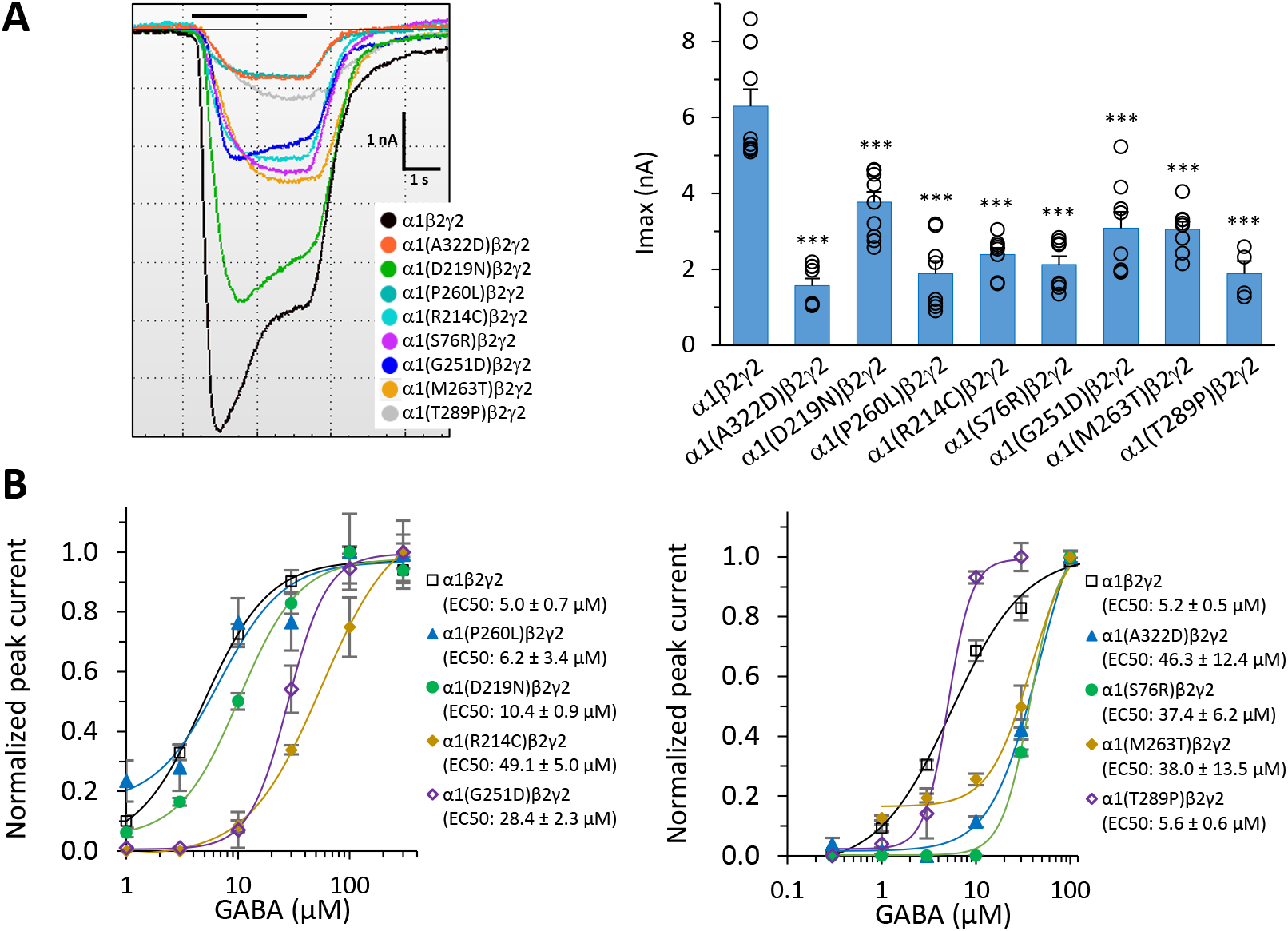
Whole-cell patch-clamp recording of selected eight GABA_A_ receptor α1 subunit DAVs. **a** HEK293T cells were transiently transfected with α1 (wild type or the indicated variants), β2, and γ2 cDNAs of GABA_A_ receptors. Forty-eight hours post transfection, whole-cell currents were recorded using the IonFlux Mercury 16 ensemble plates at a holding voltage of −60 mV. Representative whole-cell voltage-clamp recording traces are shown. Application of GABA (100 μM, 3 s) is indicated by the horizontal bar above the current traces. Quantification of the peak currents (I_max_) is shown on the right (n = 4 to 10). nA: nano Ampere. **b** Dose-response curves of GABA are shown for the calculation of EC_50_ values for wild type and the indicated α1 DAVs (n = 3 to 4). Each data point is reported as mean ± SEM. One-way ANOVA followed by post-hoc Tukey test was used for statistical analysis. ***, *p* < 0.001.

In addition, we carried out dose-response analysis to evaluate the effect of the eight DAVs on the potency of GABA, measured by the EC_50_ values (the half maximal effective concentrations) of GABA in HEK293T cells (**Fig. 2b**). Wild type α1β2γ2 receptors exhibited an EC_50_ of 5.0 ± 0.7 μM, consistent with previous reports ^15, 40^. Compared to wild type receptors, the most dramatic increases in EC_50_ values were observed in the variants of S76R (37.4 ± 6.2 μM), R214C (49.1 ± 5.0 μM), G251D (28.4 ± 2.3 μM), M263T (38.0 ± 13.5 μM), and A322D (46.3 ± 12.4 μM) (**Fig. 2b**), indicating that these variants substantially decreased GABA’s potency, consistent with their loss-of-function effects.

The above data clearly demonstrated that all of the eight selected α1 DAVs led to their loss of function according to electrophysiological experiments. Importantly, seven out of eight DAVs resulted in significantly reduced surface trafficking of GABA_A_ receptors, presumably due to protein misfolding of such variants. Therefore, correcting the folding defects of such variants to restore their surface expression has the potential to restore their functions on the plasma membrane.

### Screening a series of GABA_A_ receptor-specific modulators identifies pharmacological chaperones that increase the total protein level of **α**1 DAVs

To identify pharmacological chaperones (PCs) for α1 DAVs, we screened a series of GABA_A_ receptor modulators, including agonists, antagonists, and allosteric modulators, which are known to bind GABA_A_ receptors, namely GABA, Midazolam, Florazepam, Chlormezanone, Hispidulin, CL218872, ZK93423, Bretazenil, CGS20625, and TP003, for their capacity to increase the total protein level of α1 DAVs in HEK293T cells ^29, 46, 47^. Three stable cell lines expressing α1(D219N)β2γ2, α1(G251D)β2γ2, or α1(P260L)β2γ2 receptors were treated with these modulators at 10 μM for 24 h. Western blot analysis demonstrated that Hispidulin, TP003, and Midazolam increased total protein levels for all three α1 DAVs, and CL218872 increased total α1 protein levels for α1(D219N) and α1(P260L) significantly (**Supplementary Fig. S3**). In addition, these GABA_A_ receptor-specific modulators had no apparent effects on the total protein levels of BiP, a Hsp70 family chaperone in the ER lumen (**Supplementary Fig. S3**), indicating that they did not activate the ER stress response. All four effective compounds, namely Hispidulin ^48^, Midazolam ^49^, TP003 ^50^, and CL218872 ^51^, belong to positive allosteric modulators (PAMs), acting primarily on the benzodiazepine site at the α1+/γ2− interface (**Supplementary Fig. S1b**), and cross the blood-brain barrier. Since Hispidulin and TP003 are effective in increasing total protein levels for all three α1 DAVs, we chose them for further studies. Hispidulin is a naturally occurring flavone ^52^, whereas TP003 is an imidazopyridine that was chemically synthesized ^53^ (see **Fig. 3a** for their chemical structures).

**Fig 3.**
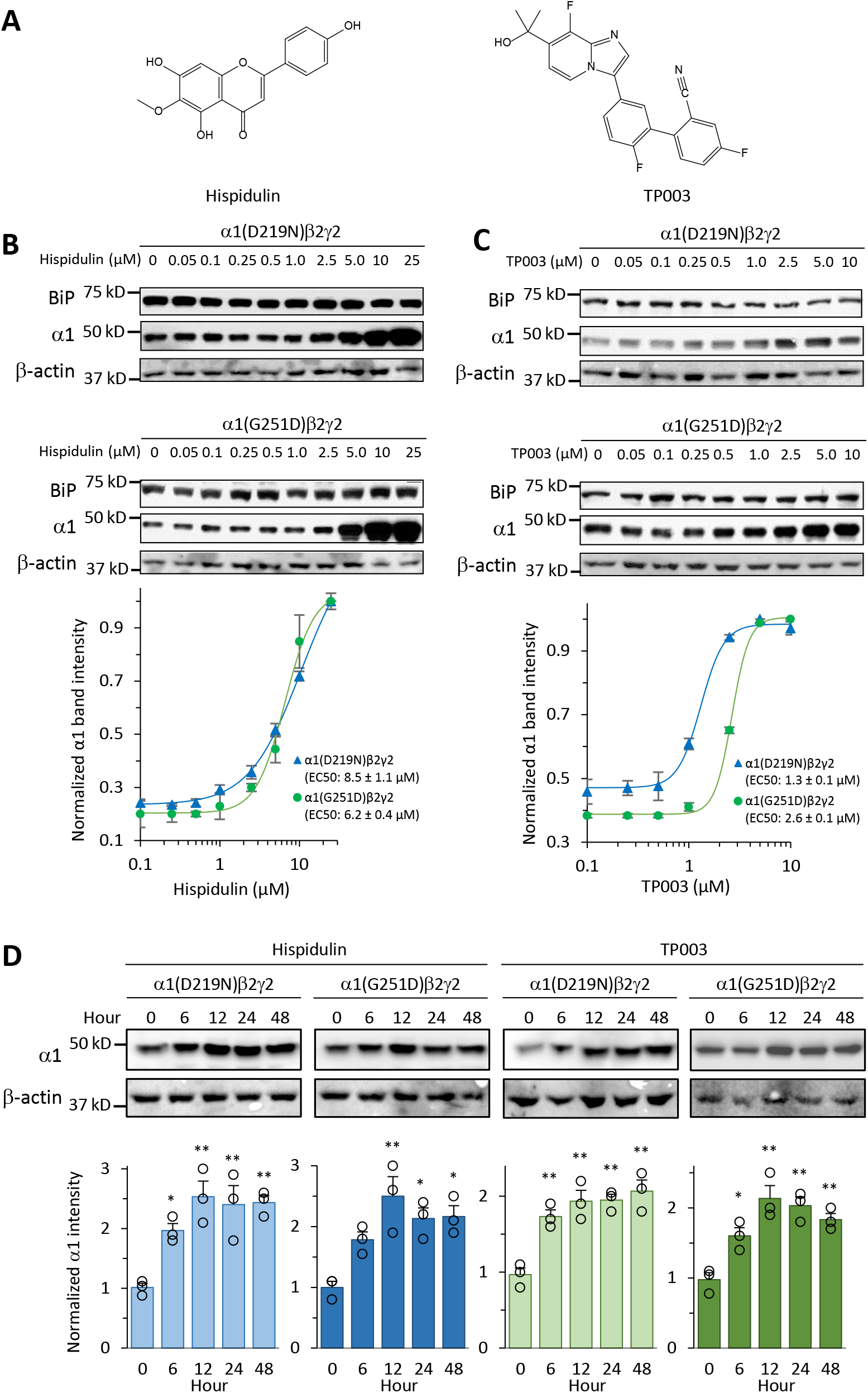
Hispidulin and TP003, positive allosteric modulators (PAMs) of GABA_A_ receptors, increase the total protein levels of α1 DAVs. **a** Chemical structures of Hispidulin and TP003. **b**, **c** Dose-response analysis of Hispidulin (**b**) and TP003 (**c**). HEK293T cells stably expressing α1(D219N)β2γ2 or α1(G251D)β2γ2 receptors were treated with DMSO vehicle control, Hispidulin (0.05 μM to 25 μM), or TP003 (0.05 μM to 10 μM) for 24h. Cells were lysed, and total proteins were subjected to Western blot analysis. EC_50_ values were calculated from the dose-response curve fitting, shown on the bottom panels (n=3). **d** Time-course analysis of Hispidulin and TP003. HEK293T cells stably expressing α1(D219N)β2γ2 or α1(G251D)β2γ2 receptors were treated with DMSO vehicle control, Hispidulin (10 μM), or TP003 (5 μM) for the indicated time. Cells were lysed, and total proteins were subjected to Western blot analysis. Quantification of the α1 band intensities was shown on the bottom panels (n=3). Each data point is reported as mean ± SEM. One-way ANOVA followed by post-hoc Tukey test was used for statistical analysis. *, *p* < 0.05; **, *p* < 0.01. Also see Supplementary Fig. S3 and Fig. S4.

To determine the optimal concentrations of Hispidulin and TP003, we carried out dose response experiments in HEK293T cells stably expressing α1(D219N)β2γ2 or α1(G251D)β2γ2 receptors. Hispidulin exhibited EC_50_ values of 8.5 ± 1.1 μM and 6.2 ± 0.4 μM for increasing α1(D219N) and α1(G251D) protein levels, respectively (**Fig. 3b**), and TP003 exhibited EC_50_ values of 1.3 ± 0.1 μM and 2.6 ± 0.1 μM for increasing α1(D219N) and α1(G251D) protein levels respectively (**Fig. 3c**). Importantly, Hispidulin and TP003 had minimal effects on wild type α1 protein levels in HEK293T cells stably expressing α1β2γ2 receptors (**Supplementary Fig. S4**), indicating that these two PCs selectively target misfolding-prone variants over wild type α1 protein. In addition, BiP protein levels remained unchanged after the application of Hispidulin (0.05 μM to 25 μM) and TP003 (0.05 μM to 10 μM) (**Fig. 3b**, **Fig. 3c**, **Supplementary Fig. S4**), indicating that Hispidulin and TP003 did not induce ER stress. Time-course experiments showed that after a single dose application of Hispidulin (10 μM) or TP003 (5 μM), total protein level increases for α1(D219N) and α1(G251D) were observed as early as 6 hours, peaked at 12-24 hours, and lasted at least to 48 hours (**Fig. 3d**). Therefore, in subsequent experiments we incubated cells for 24 h in a 10 µM concentration of Hispidulin and 5 µM of TP003, unless otherwise indicated.

### Hispidulin and TP003 promote functional surface expression of **α**1 DAVs

Because GABA_A_ receptors need to be trafficked to the plasma membrane where they act as chloride channels, we determined whether PC treatment restored the surface trafficking of misfolding-prone α1 DAVs. Cell surface biotinylation assays demonstrated that Hispidulin treatment increased the surface expression of α1(D219N) by 1.8-fold, α1(G251D) by 3.0-fold, and α1(P260L) by 2.4-fold in HEK293T cells (**Fig. 4a-c**); TP003 treatment increased the surface expression of α1(G251D) by 2.9-fold and α1(P260L) by 2.0-fold in HEK293T cells (**Fig. 4b, 4c**).

**Fig 4.**
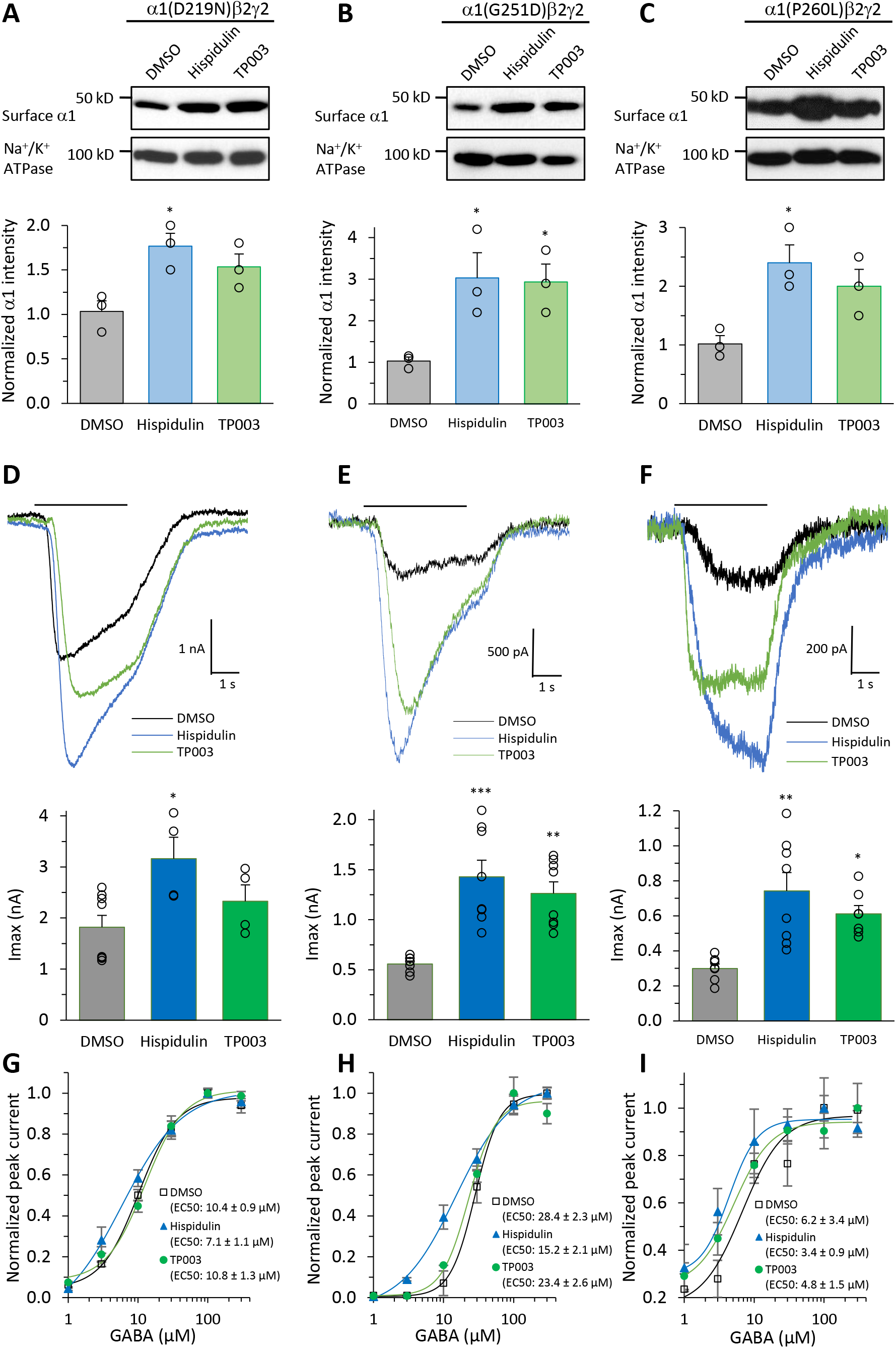
Hispidulin and TP003 promote the functional surface expression of α1 DAVs. **a-c** Effect of Hispidulin (10 µM, 24 h) or TP003 (5 µM, 24 h) on the surface protein expression of the α1 variants in HEK293T cells stably expressing α1(D219N)β2γ2 (**a**), α1(G251D)β2γ2 (**b**), or α1(P260L)β2γ2 GABA_A_ receptors (**c**) according to surface biotinylation analysis. Na^+^/K^+^ ATPase serves as a plasma membrane protein loading control. Quantification of the surface α1 band intensities was shown on the bottom panels (*n* = 3). **d-f** Effect of Hispidulin (10 µM, 24 h) or TP003 (5 µM, 24 h) on GABA-induced peak current amplitudes in HEK293T cells stably expressing α1(D219N)β2γ2 (**d**), α1(G251D)β2γ2 (**e**), or α1(P260L)β2γ2 GABA_A_ receptors (**f**). Representative whole-cell voltage-clamp recording traces are shown. The holding voltage was set at −60 mV. Application of GABA (100 μM, 3 s) is indicated by the horizontal bar above the current traces. Quantification of the peak currents (I_max_) are shown on the bottom panels (n = 4 to 8). pA: pico Ampere; nA: nano Ampere. **g-i** Dose-response curves of GABA are shown for the evaluation of the effect of Hispidulin (10 µM, 24 h) or TP003 (5 µM, 24 h) on α1(D219N)β2γ2 (**g**), α1(G251D)β2γ2 (**h**), or α1(P260L)β2γ2 GABA_A_ receptors (**i**) (n = 3 to 4). Each data point is reported as mean ± SEM. One-way ANOVA followed by post-hoc Tukey test was used for statistical analysis. * *p* < 0.05; ** *p* < 0.01; *** *p* < 0.001.

Furthermore, to determine whether the increased surface α1 DAVs form functional receptors on the plasma membrane, we used whole-cell patch-clamp electrophysiology to record GABA-induced currents. Hispidulin treatment (10 µM, 24h) significantly increased the peak current amplitudes in HEK293T cells stably expressing α1(D219N)β2γ2 receptors (from 1.8 nA to 3.2 nA) (**Fig. 4d**), α1(G251D)β2γ2 receptors (from 560 pA to 1.4 nA) (**Fig. 4e**), and α1(P260L)β2γ2 receptors (from 300 pA to 740 pA) (**Fig. 4f**). TP003 treatment (5 µM, 24h) significantly increased the peak current amplitudes in HEK293T cells stably expressing α1(G251D)β2γ2 receptors (from 560 pA to 1.3 nA) (**Fig. 4e**) and α1(P260L)β2γ2 receptors (from 300 pA to 610 pA) (**Fig. 4f**). Furthermore, we determined the effect of PC treatment on the potency of GABA. Hispidulin treatment (10 µM, 24h) reduced EC_50_ values of GABA in HEK293T cells stably expressing α1(D219N)β2γ2 receptors (from 10.4 ± 0.9 μM to 7.1 ± 1.1 μM) (**Fig. 4g**), α1(G251D)β2γ2 receptors (from 28.4 ± 2.3 μM to 15.2 ± 2.1 μM) (**Fig. 4h**), and α1(P260L)β2γ2 receptors (from 6.2 ± 3.4 μM to 3.4 ± 0.9 μM) (**Fig. 4i**), indicating that Hispidulin treatment increased the potency of GABA in these variants; however, TP003 treatment (5 µM, 24h) had minimal effects on the EC_50_ values of GABA in these variants (**Fig. 4g-i**), indicating the subtle difference between Hispidulin and TP003 on GABA_A_ receptor function possibly due to that TP003 is a weaker PAM compared to Hispidulin.

These data clearly demonstrated that Hispidulin and TP003 are capable of restoring the surface trafficking and whole-cell currents of GABA_A_ receptors by incorporating α1 DAVs into functional receptors. Since Hispidulin and TP003 directly bind GABA_A_ receptors at the benzodiazepine sites, they are promising PCs for further clinical development to rescue misfolding-prone GABA_A_ receptor variants.

### Hispidulin and TP003 promote the folding, assembly, and anterograde trafficking of the α1(G251D) variant

We next determined the mechanism of action of Hispidulin and TP003 in correcting the function of α1(G251D) variant. Presumably, binding of Hispidulin and TP003 to the α1(G251D) variant improves stability and thus enhances the assembly and forward trafficking of GABA_A_ receptors. We used Förster resonance energy transfer (FRET) assay to quantify the subunit-subunit assembly in HEK293T cells ^17^. CFP-tagged α1(G251D) and YFP-tagged β2 exhibited a FRET efficiency of 21.4 ± 1.4%; Hispidulin or TP003 treatment significantly increased the FRET efficiency to 42.5 ± 1.2% and 41.0 ± 2.1%, respectively (**Fig. 5a, 5b**), indicating that PC treatments enhanced the assembly between α1(G251D) and β2 subunits to form mature pentameric receptors.

**Fig 5.**
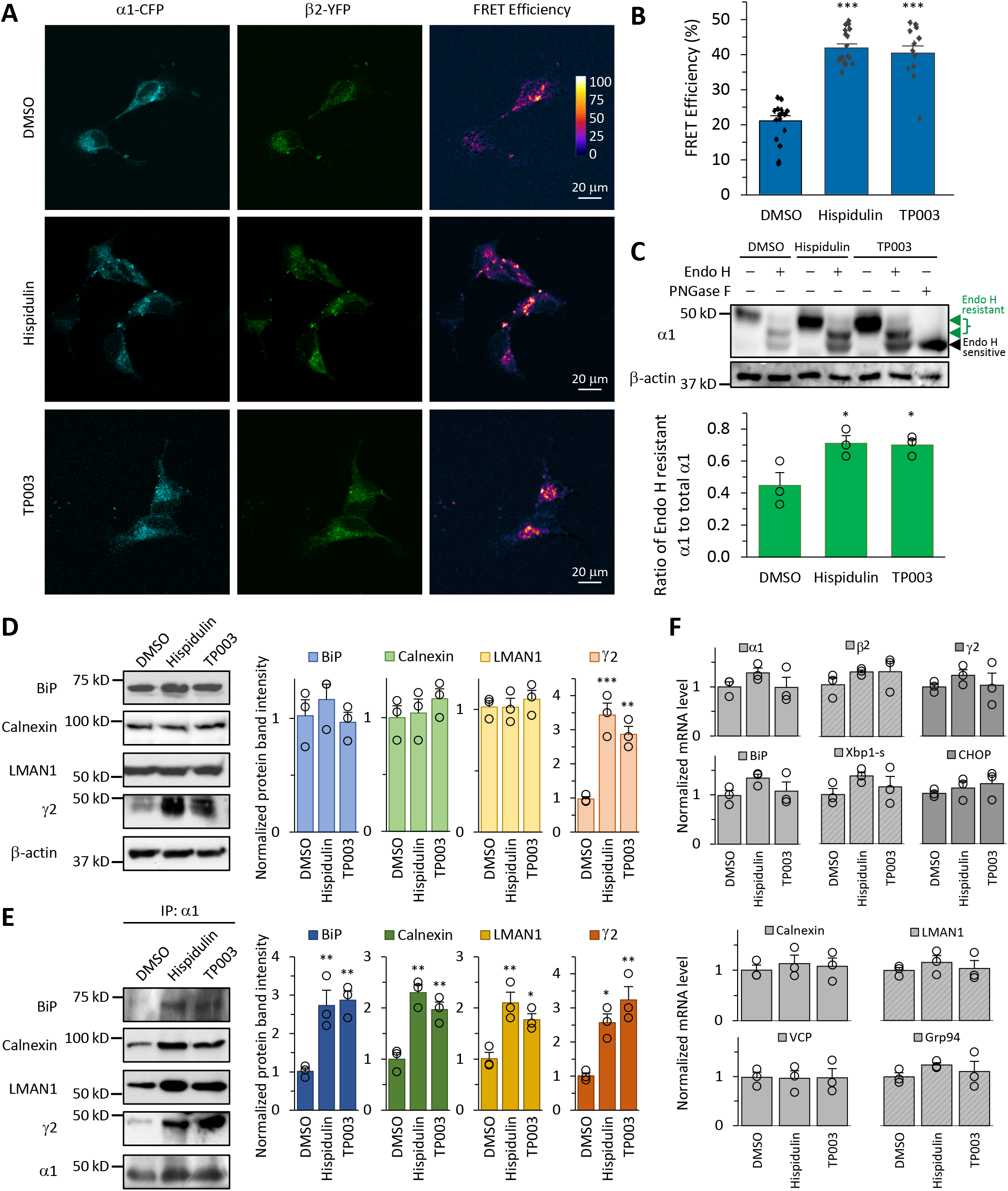
Hispidulin and TP003 promote the folding, assembly, and forward trafficking of misfolding-prone α1(G251D) variant. **a,b** Hispidulin (10 µM, 24 h) or TP003 (5 µM, 24 h) increases the FRET efficiency between CFP-tagged α1(G251D) subunit and YFP-tagged β2 subunit of GABA_A_ receptors. HEK293T cells were transfected with CFP-tagged α1(G251D) subunit, YFP-tagged β2 subunit, and γ2 subunit; Twenty-four hours post transfection, cells were treated with DMSO, Hispidulin (10 µM) or TP003 (5 µM). Forty-eight hours post transfection, pixel-based FRET was used to calculate the FRET efficiency between α1(G251D)-CFP and β2-YFP. Representative images were shown for the CFP channel (1st columns), YFP channel (2nd columns), and FRET efficiency (3rd columns) (**a**). Scale bar = 20 μm. FRET efficiency was calculated from 30-40 cells from at least three transfections using the ImageJ PixFRET plug-in, shown in **b**. **c** Hispidulin (10 µM, 24 h) or TP003 (5 µM, 24 h) treatment increases the ER-to-Golgi trafficking efficiency of α1(G251D) in HEK293T cells stably expressing α1(G251D)β2γ2 GABA_A_ receptors. The Peptide-N-Glycosidase F (PNGase F) enzyme cleavage serves as a control for unglycosylated α1 subunits (lane 7). After endo H digestion, α1 subunits with a molecular weight that is equal to unglycosylated α1 were labeled as endo H-sensitive, whereas those with higher molecular weights were labeled as endo H-resistant. Quantification of the ratio of endo H-resistant α1 / total α1 subunit bands, as a measure of the ER-to-Golgi trafficking efficiency, is shown on the bottom (n=3). **d** Effect of Hispidulin (10 µM, 24 h) and TP003 (5 µM, 24 h) on the protein levels of selected chaperones and γ2 subunits in HEK293T cells stably expressing α1(G251D)β2γ2 receptors (n=3). **e** Effect of Hispidulin (10 µM, 24 h) and TP003 (5 µM, 24 h) on the interactions between α1(G251D) and selected chaperones or γ2 in HEK293T cells stably expressing α1(G251D)β2γ2 receptors. Quantification of the ratio of proteins of interest / α1(G251D) post immunoprecipitation is shown in the right panels (n=3). IP: immunoprecipitation. **f** Effect of Hispidulin (10 µM, 24 h) and TP003 (5 µM, 24 h) on the mRNA levels of genes of interest in HEK293T cells stably expressing α1(G251D)β2γ2 receptors, assessed by quantitative RT-PCR (n=3). Each data point is reported as mean ± SEM. One-way ANOVA followed by post-hoc Tukey test was used for statistical analysis. * *p* < 0.05; ** *p* < 0.01; *** *p* < 0.001. Also see **Supplementary Table S2**.

Furthermore, to determine whether the enhanced assembly resulted in enhanced anterograde trafficking after drug treatment, we used an endoglycosidase H (endo H) enzyme digestion assay to quantify the ER-to-Golgi trafficking efficiency. The endo H enzyme selectively removes the N-linked glycans in the ER, but not in the Golgi after the remodeling of this oligosaccharide chain in the Golgi. Therefore, properly folded and assembled α1 glycoforms that can traffic to the Golgi are resistant to endo H digestion. Since the α1 subunit has two glycosylation sites (Asn38 and Asn138) in the ER, endo H digestion produced two endo H-resistant bands (**Fig. 5c**, lanes 2, 4, and 6). Endo H-sensitive bands have the same apparent molecular weight as unglycosylated α1 subunits, as generated by PNGase F enzyme digestion (**Fig. 5c**, lane 7). Hispidulin or TP003 treatment clearly increased the upper two endo H-resistant α1(G251D) bands in HEK293T cells (**Fig. 5c**, cf. lanes 4 and 6 to lane 2), indicating that PC treatment increased post-ER α1(G251D) glycoforms. Moreover, Hispidulin or TP003 treatment significantly increased the ratio of endo H resistant α1 / total α1 (**Fig 5c**, cf. lanes 4 and 6 to lane 2, quantification shown on the bottom), indicating that PC treatment enhanced the trafficking efficiency of the α1(G251D) variant from the ER to the Golgi.

Since the ER proteostasis network orchestrates the folding, assembly, degradation, and trafficking of GABA_A_ receptors ^13^, we determined the effect of Hispidulin or TP003 on adapting this network. Previously, we demonstrated that ER resident chaperones, including BiP and calnexin, positively regulate the productive folding of GABA_A_ receptors ^15, 16^, and LMAN1 (aka ERGIC53) promotes their anterograde trafficking ^18^. Although Hispidulin or TP003 treatment did not change the protein levels of these chaperones (**Fig. 5d**), co-immunoprecipitation assay demonstrated that they significantly enhanced the interactions between α1(G251D) and BiP/calnexin/LMAN1 (**Fig. 5e**). Such enhanced interactions after PC treatment would shift the partition of the variant from the ER to engage the trafficking machinery for their transport to the Golgi and onward to the surface. In addition, Hispidulin or TP003 treatment substantially increased the interaction between α1(G251D) and γ2 subunits of GABA_A_ receptors (**Fig. 5e**), indicating that indeed PC treatment promoted the assembly of GABA_A_ receptors, consistent with the FRET results (**Fig. 5a**).

Furthermore, quantitative RT-PCR demonstrated that Hispidulin or TP003 treatment did not change the mRNA levels of α1(G251D), β2, and γ2 subunits of GABA_A_ receptors (**Fig. 5f**), indicating the post-transcriptional effect of PC treatment on rescuing α1 DAVs. Since we previously reported that the genetic or pharmacologic activation of the unfolded protein response (UPR) is capable of restoring the folding, trafficking, and function of epilepsy-associated GABA_A_ receptor variants ^25, 26^, we tested whether Hispidulin or TP003 induced the UPR. Quantitative RT-PCR revealed that Hispidulin or TP003 treatment did not change the transcription of BiP, spliced XBP1, and CHOP, which are markers for the activation of the three established arms of the UPR, namely ATF6, IRE1, and PERK ^54^ (**Fig. 5f**), indicating that PC treatment did not activate the UPR. Consistently, the transcription of the UPR downstream targets, including calnexin, LMAN1, VCP, and Grp94 ^55^, was not changed after PC treatment (**Fig. 5f**).

### Hispidulin and TP003 reduce the degradation of the **α**1(G251D) variant

We next evaluated whether Hispidulin or TP003 treatment inhibited the degradation of the α1(G251D) variant due to their stabilization effect on GABA_A_ receptors. Cycloheximide, a potent protein synthesis inhibitor, was applied to HEK293T cells stably expressing α1(G251D)β2γ2 receptors to determine the degradation kinetics of α1(G251D) protein. Cycloheximide-chase assay demonstrated that Hispidulin or TP003 treatment significantly increased the remaining protein levels of α1(G251D) at 0.5 h and at 1 h post the application of cycloheximide (**Fig. 6a**), indicating that both compounds attenuated the degradation of the misfolding-prone α1(G251D) variant.

**Fig 6.**
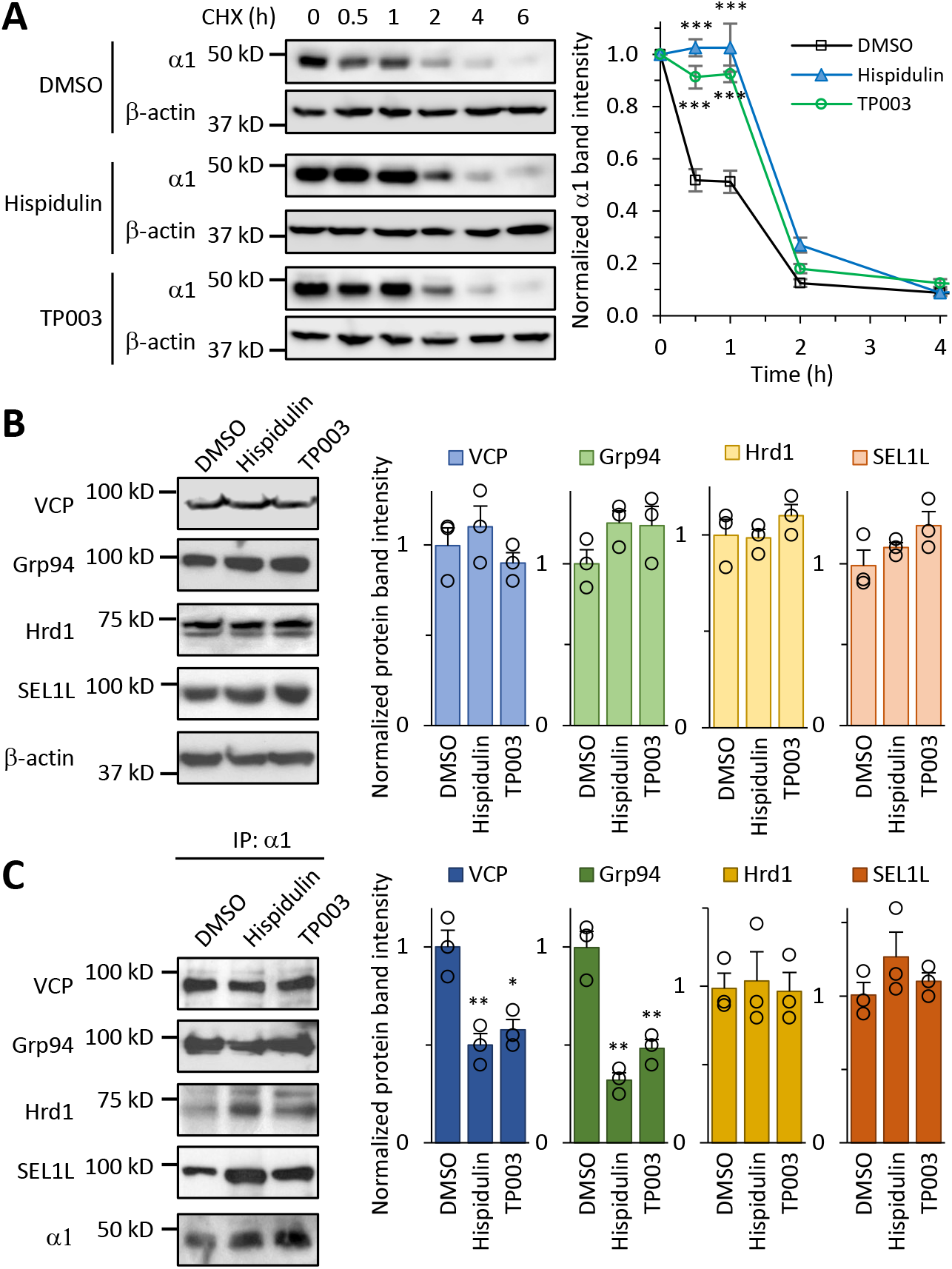
Hispidulin and TP003 inhibit the ERAD of the misfolding-prone α1(G251D) variant. **a** HEK293T cells stably expressing α1(G251D)β2γ2 receptors were treated with DMSO, Hispidulin (10 µM), and TP003 (5 µM) for 24 h. Then cycloheximide (100 μg / mL) (CHX), a potent protein synthesis inhibitor, was added to the cell culture media for the indicated time before cell lysis and Western blot analysis. Quantification of the normalized remaining α1 protein band intensity was shown on the right (n = 3). **b** Effect of Hispidulin (10 µM, 24 h) and TP003 (5 µM, 24 h) on the protein levels of selected ERAD factors in HEK293T cells stably expressing α1(G251D)β2γ2 receptors (n=3). **c** Effect of Hispidulin (10 µM, 24 h) and TP003 (5 µM, 24 h) on the interactions between α1(G251D) and selected ERAD factors in HEK293T cells stably expressing α1(G251D)β2γ2 receptors. Quantification of the ratio of proteins of interest / α1(G251D) post immunoprecipitation is shown in the right panels (n=3). IP: immunoprecipitation. Each data point is reported as mean ± SEM. ANOVA followed by post-hoc Tukey test was used to evaluate the statistical significance. *, *p* < 0.05; **, *p* < 0.01; ***, *p* < 0.001.

We continued to analyze the effect of PC treatments on ERAD factors to reduce α1(G251D) degradation. Previously, we demonstrated that Grp94, an ER lumenal Hsp90 chaperone, recognizes misfolded α1 subunits for their targeting to the ubiquitin E3 ligase Hrd1/Sel1L complex; then, VCP, an ATPase, extracts ubiquitinated α1 subunits from the ER membrane to the cytosol for proteasomal degradation ^12, 56^. Western blot analysis revealed that Hispidulin or TP003 treatment did not change the protein levels of all tested ERAD factors, including Grp94, Hrd1, Sel1L, and VCP (**Fig. 6b**). However, co-immunoprecipitation assay demonstrated that PC treatments significantly reduced the interactions between the α1(G251D) variant and Grp94/VCP (**Fig. 6c**), indicating that both compounds inhibited the Grp94/VCP-mediated ERAD pathway of the misfolding-prone α1(G251D) variant.

In summary, our data clearly showed that Hispidulin and TP003 both reduced the degradation and promoted the productive folding, assembly, and forward trafficking of the α1 DAVs, without activation of the UPR. As a result, PC treatments restored the function of the variant-containing receptors that reach the plasma membrane.

### Hispidulin and TP003 restore surface trafficking of the **α**1(G251D) variant in human induced pluripotent stem cell (iPSC)-derived neurons by adapting the ER proteostasis network

To mimic the native neuronal environment of GABA_A_ receptor variants, we developed isogenic human iPSCs carrying the heterozygous α1(G251D) variant, one of the variants that we selected, using CRISPR/Cas9 genome-editing (**Fig. 7a**, **Supplementary Fig. S5**). To induce the efficient differentiation of iPSCs into a homogenous population of GABAergic neurons, we expressed neurogenic transcription factors, including ASCL1 and DXL2, under the control of a doxycycline-induced promoter. The transduced iPSCs treated with deoxycycline underwent rapid neurogenesis in about 1 week ^57^. The induced GABAergic neurons (iNs) were further matured for about 3 weeks before imaging experiments were carried out.

**Fig 7.**
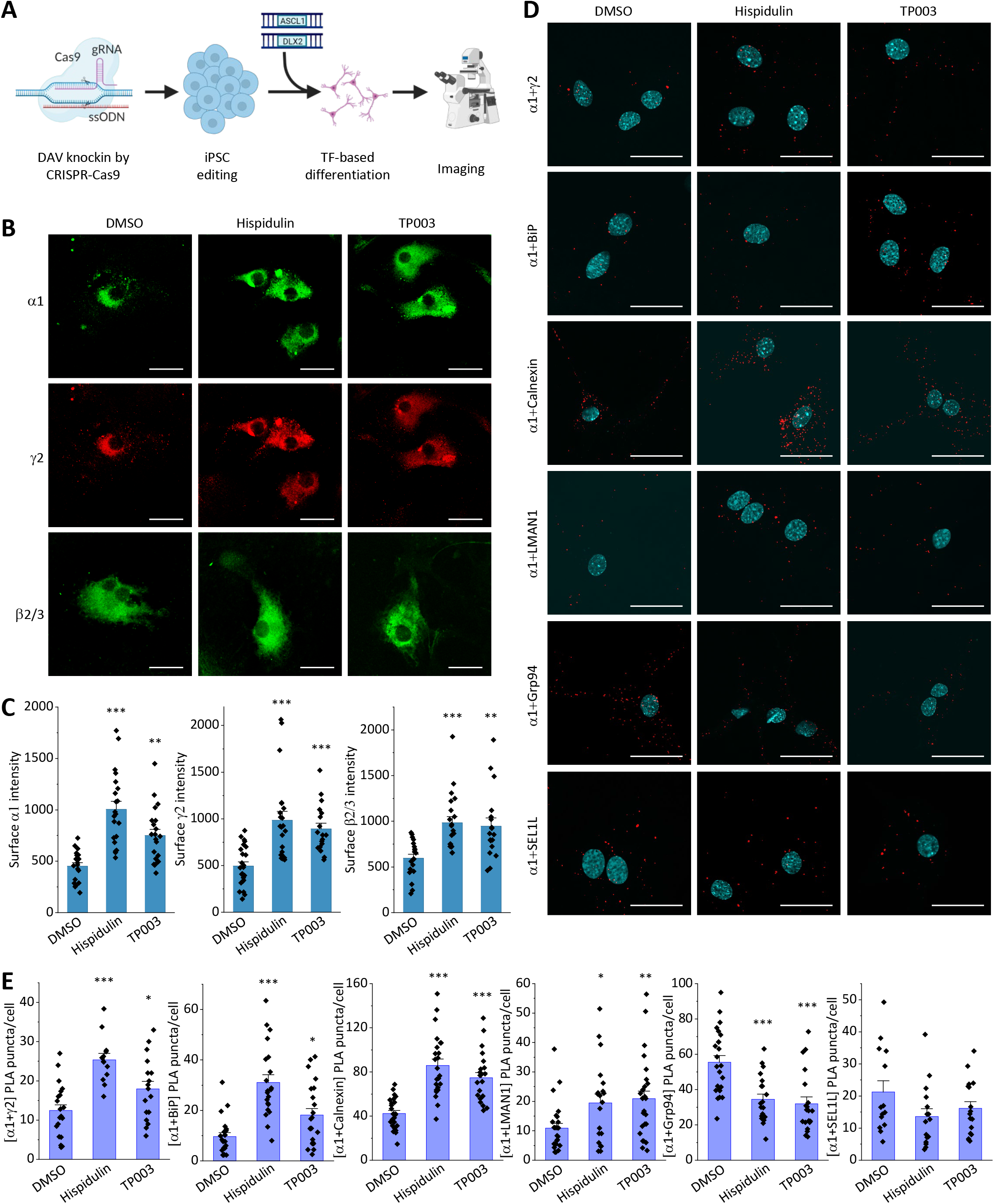
Hispidulin and TP003 promotes the surface trafficking of the α1(G251D) variant in hiPSC-derived GABAergic neurons. **a** Schematic of the generation of α1(G215D) knockin in hiPSCs and the transcription factor (TF)-based differentiation into GABAergic neurons. **b** Effect of Hispidulin (10 µM, 24 h) and TP003 (5 µM, 24 h) on the surface expression of GABA_A_ receptor subunits in hiPSC-derived GABAergic neurons carrying the α1(G215D) variant. Surface GABA_A_ receptors were stained using anti-α1 subunit, anti-β2/β3 subunit, or anti-γ2 subunit antibodies without membrane permeabilization. 50-75 neurons from at least three differentiations were imaged by confocal microscopy for each condition. Scale bar = 20 μm. **c** Quantification of the fluorescence intensity of the surface GABA_A_ receptor subunits after background correction. **d** Effect of Hispidulin (10 µM, 24 h) and TP003 (5 µM, 24 h) on the interactions between α1(G251D) and selected ER proteostasis network components. PLA was used to determine the endogenous protein-protein interactions. 35-70 neurons from at least three differentiations were imaged by confocal microscopy for each condition. Scale bar = 20 μm. **e** Quantification of the PLA puncta number per cell was achieved using the ImageJ Analyze Particles plug-in. Each data point is reported as mean ± SEM. ANOVA followed by post-hoc Tukey test was used to evaluate the statistical significance. *, *p* < 0.05; **, *p* < 0.01; ***, *p* < 0.001. Also see **Supplementary Fig. S5**.

Since insufficient presence of GABA_A_ receptors at the synaptic sites resulting from the pathogenic α1(G251D) variant causes epilepsy, we explored whether PC treatment enhanced the surface trafficking of major synaptic GABA_A_ receptor subunits (α1, β2/β3, and γ2) in iNs carrying the α1(G251D) variant. Immunofluorescence staining of the surface proteins clearly demonstrated that Hispidulin (10 μM, 24 h) or TP003 (5 μM, 24h) treatment increased the surface fluorescence intensities for the major synaptic GABA_A_ subunits that form pentameric receptors (**Fig. 7b, 7c**). The enhanced surface presence of GABA_A_ receptors would likely correct the impaired GABAergic input.

Furthermore, we investigated how PC treatment adapted the ER proteostasis network to restore the surface trafficking of α1(G251D) in the human disease-relevant iN cellular environment. *In situ* Proximity Ligation Assay (PLA) was used to determine the endogenous interactions between α1(G251D) and ER proteostasis network components in iNs (**Fig. 7d**). Quantification of the PLA puncta was achieved using the ImageJ software with the built-in macro Analyze Particles (**Fig. 7e**). PLA showed that Hispidulin (10 μM, 24 h) or TP003 (5 μM, 24h) treatment increased the interactions between α1(G251D) and γ2 subunits (**Fig. 7d**, row 1, **Fig. 7e**), indicating that PC treatment enhanced the assembly of GABA_A_ receptor subunits to form pentameric receptors. In addition, Hispidulin or TP003 treatment increased the interactions between α1(G251D) and pro-folding chaperones, including BiP and calnexin (**Fig. 7d**, rows 2 and 3, **Fig. 7e**), as well as a trafficking receptor LMAN1 (**Fig. 7d**, row 4, **Fig. 7e**). In contrast, PC treatment reduced the interactions between α1(G251D) and an ERAD factor Grp94 (**Fig. 7d**, row 5, **Fig. 7e**). In summary, our data in iNs provided strong evidence that PC treatment adapted the GABA_A_ receptor proteostasis network by promoting the folding, assembly, and forward trafficking of GABA_A_ receptors as well as reducing their ERAD. As a result, PC treatment significantly enhanced the surface trafficking of α1 DAVs in iNs.

## Discussion

Here, we characterized proteostasis deficiency of selected DAVs of GABA_A_ receptor α1 subunits. Among the known > 200 variants, we chose eight pathogenic variants for our study that cover a broad phenotypic spectrum, including severe epileptic encephalopathies, such as Dravet syndrome, and milder IGE, such as JME ^58^. Additionally, the selected DAVs occupy the NTD and TM helices, representing a diverse spatial distribution in the tertiary protein structure of GABA_A_ receptors. Clearly, all eight α1 DAVs led to reduced peak current amplitudes in patch-clamp recordings (**Fig. 2a**). Such loss of function of GABA_A_ receptors due to these DAVs is consistent with the idea that reduced GABAergic inhibition in the central nervous system causes the hyperexcitability tendency and thus neurodevelopment disorders, such as epilepsy ^59^. Furthermore, seven out of eight DAVs decreased their surface expression, suggesting that trafficking deficiency resulting from protein conformational defects is the major disease-causing mechanism for GABA_A_ receptor variants. Certainly, such a claim would be fully supported by a comprehensive characterization of all DAVs, calling for the need of the development of high throughput assay to quantify the receptor surface expression. Nonetheless, this study provides strong evidence toward that end. In addition, prediction of pathogenic probability of the α1 subunit suggests several regions that are intolerant to variations, such as TM1-3 and the NTD loop that is close to TM1.

To correct the function of α1 DAVs, we identified Hispidulin and TP003 as effective PCs that restored the surface trafficking and GABA-induced currents for α1(D219N), α1(G251D), and α1(P260L), without a substantial effect on wild type receptors. Hispidulin and TP003, as well as other effective modulators Midazolam and CL218872, bind to the benzodiazepine site of pentameric GABA_A_ receptors. The canonical high-affinity benzodiazepine site is located in the NTD at the α1+/γ2− interface (**Supplementary Fig. S1b**); in addition, recent structural studies identified low-affinity benzodiazepine sites in the transmembrane domains at the β+/α1− interfaces ^60^. Hispidulin and TP003 are membrane permeable and cross the blood-brain barrier ^48, 50^, and presumably, they will enter the ER to interact with GABA_A_ receptor subunits since ER membrane is more permeable than the plasma membrane ^61^. PC binding to GABA_A_ receptor subunits in the ER stabilizes the otherwise degraded GABA_A_ variants and promotes their assembly and anterograde trafficking from the ER to the Golgi and onward to the plasma membrane (**Fig. 8**), thus restoring GABA-induced currents on the plasma membrane. PC treatment adapts the ER proteostasis by adjusting the interactions between DAVs and GABA_A_ receptor proteostasis network (**Fig. 5–7**) ^13^: the interactions between the variants and pro-folding chaperones (BiP and calnexin) and a trafficking factor (LMAN1) are enhanced, whereas the interactions between the variants and ERAD factors (Grp94 and VCP) are reduced by PC chaperone treatment (**Fig 8**). Since Hispidulin and TP003 are PAMs of GABA_A_ receptors, in addition to their PC effects on restoring the trafficking of GABA_A_ variants, presumably, their application could potentially increase GABA-induced currents further once the receptors are able to reach the plasma membrane, adding the benefits of using these compounds to correct the function of GABA_A_ variants.

**Fig 8.**
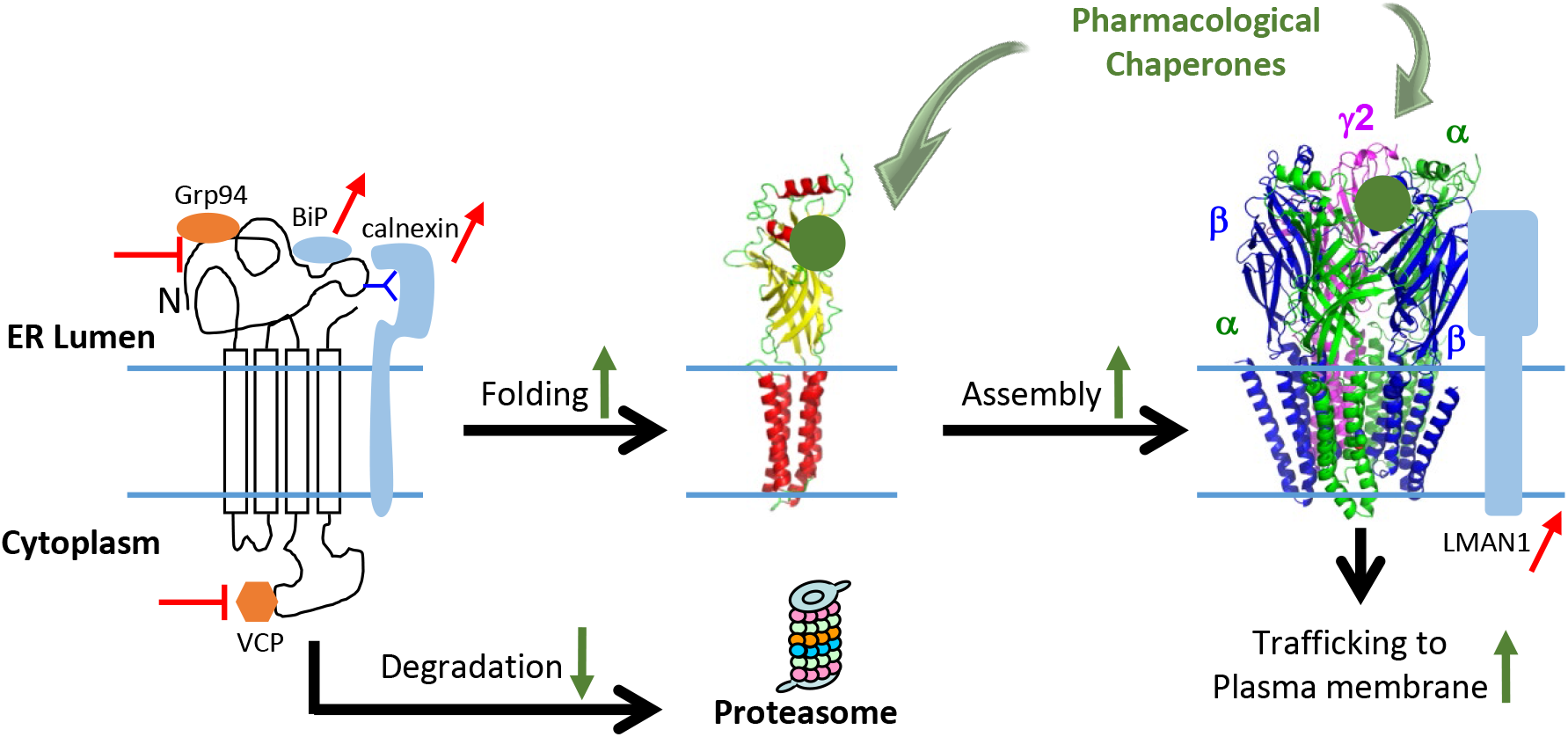
Proposed mechanism of action of Hispidulin and TP003 on rescuing α1 DAV trafficking. Hispidulin and TP003 act as pharmacological chaperones to bind GABA_A_ receptor variants to stabilize them in the ER. Pharmacological chaperone treatment enhances the assembly of the variants and their forward trafficking to the Golgi and plasma membrane. The effect of Hispidulin and TP003 is also reflected from the enhanced folding and reduced degradation of the variants. Consistently, the interactions between the variants and pro-folding chaperones (BiP and calnexin) and a trafficking factor (LMAN1) are enhanced, whereas the interactions between the variants and ERAD factors (Grp94 and VCP) are reduced by pharmacological chaperone treatment.

Hispidulin is a promising candidate to be further developed as a therapeutic agent to treat genetic epilepsies since it is a naturally occurring flavone ^52, 62^; moreover, it was previously reported that application of Hispidulin at 10 mg / kg body weight per day in a Mongolian gerbil epilepsy model after seven days reduced seizures ^48^. Here, we demonstrated that Hispidulin treatment substantially increased the surface expression of α1 DAVs, including α1(D219N), α1(G251D), and α1(P260L), and thus restored their GABA-induced currents, indicating that the pharmacological chaperoning strategy is effective in correcting the function of a variety of clinical variants that cause different disease severities. Furthermore, we developed an isogenic, human iPSC model carrying the α1(G251D) variant and used induced iPSC-derived GABAergic neurons to mimic the native cellular environments for GABA_A_ receptors. We demonstrated that Hispidulin treatment substantially promoted the surface trafficking of the α1(G251D) variant in the iPSC-derived neuron model. There is an urgent need to develop new therapeutic strategy to treat genetic epilepsy resulting from loss of function of GABA_A_ receptors since many patients with GABA_A_ variants are resistant to current anti-seizure drugs ^9, 63^. Because Hispidulin effectively corrected the defects of GABA_A_ variants and it is a well-tolerated natural product that crosses the blood-brain barrier, Hispidulin holds great potential to be further developed to treat genetic epilepsies resulting from GABA_A_ receptor functional deficiency.

Previously, we identified a number of proteostasis regulators that correct the proteostasis deficiency and thus function of GABA_A_ DAVs, including HDAC inhibitors, such as SAHA ^15^, calcium channel blockers, such as verapamil ^16^, a potent BiP activator BIX ^25^, FDA-approved drugs dinoprost and dihydroergocristine ^24^, and stress-independent activators of the ATF6 arm of the UPR, including AA147 and AA263 ^26^. These proteostasis regulators remodel the ER proteostasis network by operating on chaperones and ERAD factors to stabilize the GABA_A_ variants without direct interactions between proteostasis regulators and GABA_A_ receptors: ATF6 activators AA147 and AA263, BIX, and SAHA all upregulated the BiP protein levels as a way to promote the folding of the variants, whereas dinoprost and dihydroergocristine downregulated the protein levels of certain ERAD factors, such as Hrd1 and SEL1L. The mechanism of action of proteostasis regulators is distinct from that of PCs, such as Hispidulin and TP003, since PCs can stabilize the variants through their direct interactions with GABA_A_ receptors without changing the protein levels of chaperones and ERAD factors. It is interesting that both proteostasis regulators and PCs can promote the folding and forward trafficking of GABA_A_ receptors by enhancing the interactions between chaperones/trafficking factors and GABA_A_ receptors as well as inhibiting the ERAD of GABA_A_ receptors by attenuating the interactions between ERAD factors and GABA_A_ receptors. Importantly, both proteostasis regulators and PCs can specifically correct the surface trafficking and thus function of misfolding-prone GABA_A_ variants when compared to wild type receptors. Therefore, both proteostasis regulators and PCs are promising candidates for the treatment of genetic epilepsies resulting from GABA_A_ protein conformational defects. Due to their distinct mechanism of action, it is expected that co-application of proteostasis regulators and PCs could yield additive/synergistic rescue of proteostasis deficiency of GABA_A_ variants, which deserves future investigation.

## Methods

### Reagents

GABA (catalog#: A2129), Midazolam (catalog#: M-908), and Florazepam (catalog#: F-003) were purchased from Sigma. Chlormezanone (catalog #: 50-199-8659) was obtained from Fisher Scientific. Hispidulin (catalog#: 2174), CL218872 (catalog#: 1709), ZK93423 (catalog#: 1994), Bretazenil (catalog#: 3568), CGS20625 (catalog#: 2467), and TP003 (catalog#: 4414) were purchased from Tocris Bioscience. The pCMV6 plasmids containing human GABA_A_ receptor α1 (Uniprot no. P14867-1) (catalog #: RC205390), β2 (isoform 2, Uniprot no. P47870-1) (catalog #: RC216424), γ2 (isoform 2, Uniprot no. P18507-2) (catalog #: RC209260), and pCMV6 Entry Vector plasmid (pCMV6-EV) (catalog #: PS100001) were obtained from Origene. The human GABA_A_ receptor α1 subunit missense mutations (S76R, R214C, D219N, G251D, P260L, M263T, T289P, and A322D) were constructed using QuikChange II site-directed mutagenesis Kit (Agilent Genomics, catalog #: 200523). Enhanced cyan fluorescent protein (CFP) was inserted between Lys364 and Asn365 in the TM3-TM4 intracellular loop of the α1 subunit, and enhanced yellow fluorescent proteins (YFP) was inserted between Lys359 and Met360 in the TM3-TM4 intracellular loop of the β2 subunit by using the GenBuilder cloning kit (GenScript, catalog #: L00701), as previously described ^17^. All cDNA sequences were confirmed by DNA sequencing.

### Antibodies

The mouse monoclonal anti-GABA_A_ receptor α1 subunit antibody (clone BD24, catalog #: MAB339), mouse monoclonal anti-GABA_A_ receptor β2/3 subunit antibody (clone 62-3G1, catalog #: 05-474), and rabbit polyclonal anti-γ2 subunit antibody (catalog #: AB5559) were obtained from Millipore (Burlington, MA). The mouse monoclonal anti-GABA_A_ α1 subunit antibody (catalog #: 224211), rabbit polyclonal anti-GABA_A_ α1 subunit antibody (catalog #: 224203), and rabbit polyclonal anti-GABA_A_ γ2 antibody (catalog #: 224003) were obtained from Synaptic systems. The mouse monoclonal anti-β-actin (catalog #: A1978) antibody came from Sigma (St. Louis, MO). The fluorescent anti-β-actin antibody Rhodamine came from Biorad (catalog #: 12004163). The mouse monoclonal anti-Grp78 (BiP) antibody (catalog #: AM8572b) and rabbit polyclonal anti-VCP antibody (catalog #: AP6920b) were obtained from Abgent. The rabbit polyclonal anti-BiP antibody (catalog #: ab21685), rabbit monoclonal anti-LMAN1 antibody (catalog #: ab125006), and rabbit polyclonal anti-Na^+^/K^+^ ATPase antibody (catalog #: AB76020) was obtained from Abcam (Waltham, MA). The rabbit polyclonal anti-calnexin (catalog #: ADI-SPA-860-F) and rat polyclonal anti-Grp94 (catalog #: ADI-SPA-850-F) antibodies were purchased from Enzo Life Sciences. The rabbit polyclonal anti-calnexin antibody (catalog #: 10427-2-AP), rabbit polyclonal anti-Grp94 antibody (catalog #: 14700-1-AP), and rabbit polyclonal anti-Hrd1 antibody (catalog #: 13473-1-AP) was obtained from Proteintech. The rabbit polyclonal anti-Sel1L antibody (catalog #: PA5-88333) was obtained from ThermoFisher.

### Prediction of pathogenic probability of missense variants

The Rhapsody webserver (http://rhapsody.csb.pitt.edu) was used to predict the pathogenic probability of missense variants of the human GABA_A_ receptor α1 subunit ^37^.

### Cell culture and transfection

HEK293T cells (#CRL-3216, donor sex: female) were obtained from ATCC. Cells were maintained in Dulbecco’s Modified Eagle Medium (DMEM) (Fisher Scientific, catalog #: 10-013-CV) with 10% heat-inactivated fetal bovine serum (FBS) (Fisher Scientific, catalog #: SH30396.03HI) and 1% Penicillin-Streptomycin (Fisher Scientific, catalog #: SV30010) at 37°C in 5% CO_2_. Monolayers were passaged upon reaching confluency with 0.05% trypsin protease (Fisher Scientific, catalog #: SH30236.01). Cells were grown in 6-well plates or 10-cm dishes and allowed to reach 50%-70% confluency before transient transfection using TransIT-2020 (Mirus Bio, catalog #: MIR 5400) according to the manufacturer’s instruction.

Monoclonal HEK293T cells stably expressing α1β2γ2, α1(D219N)β2γ2, α1(G251D)β2γ2, or α1(P260L)β2γ2 GABA_A_ receptors were generated using the G-418 selection and limiting dilution method, as previously described ^25^. Briefly, cells were transfected with α1:β2:γ2 (1:1:1), α1(D219N):β2:γ2 (1:1:1), α1(G251D):β2:γ2 (1:1:1) or α1(P260L):β2:γ2 (1:1:1) plasmids and selected in DMEM supplemented with 0.8 mg/mL G418 (Enzo Life Sciences) for 10 days. Cells were then diluted to 5 cells / mL, and 100 μL of cells were added into each well of 96-well plates to ensure a single cell distribution per well. Single cell colonies were inspected under a microscope and expanded. Western blot analysis was used to evaluate the expression of α1, β2, and γ2 subunits of GABA_A_ receptors. Positive monoclonal stable cells were maintained in DMEM supplemented with 0.4 mg/mL G418.

### Western blot analysis

Cells were harvested and lysed with lysis buffer (50 mM Tris, pH 7.5, 150 mM NaCl, and 2 mM *n*-Dodecyl-β-D-maltoside (DDM) (GoldBio, catalog #: DDM5)) supplemented with complete protease inhibitor cocktail (Roche, catalog #: 4693159001). Cell lysates were cleared by centrifugation (20,000 × *g*, 10 min, 4 °C): the supernatant was collected as total proteins; the pellet was washed with Dulbecco’s phosphate-buffered saline (DPBS) (Fisher Scientific, catalog #: SH3002803) and re-suspended in 2x Laemmli sample buffer containing 2% SDS (Biorad, catalog #: 1610737). Protein concentration was measured by MicroBCA assay (Thermo Scientific, catalog #: 23235). For non-reducing protein gels, total proteins were loaded in the Laemmli sample buffer; for reducing protein gels, total proteins were loaded in the Laemmli sample buffer supplemented with 5% 2-mercaptoethanol (Sigma, catalog #: M3148) to reduce the disulfide bonds. Protein samples were separated in an 8% SDS-PAGE gel. Western blot analysis was performed using appropriate antibodies. Band intensity was quantified using Image J software from the NIH.

To remove asparaginyl-*N*-acetyl-D-glucosamine in the Asn-linked glycans installed on the α1 subunit of GABA_A_ receptors in the ER, total proteins isolated from cell lysates were digested with Endoglycosidase H (endo H) enzyme (New England Biolabs, catalog #: P0703L) with G5 reaction buffer at 37°C according to manufacturer’s instruction and the published procedure ^15^. The Peptide-N-Glycosidase F (PNGase F) enzyme (New England Biolabs, catalog #: P0704L)-treated samples served as a control for unglycosylated α1 subunits. Treated samples were then subjected to Western blot analysis.

### Biotinylation of cell surface proteins

HEK293T cells stably expressing GABA_A_ receptors variants were plated in 6-cm dishes for surface biotinylation experiments according to the published procedure ^15^. Briefly, intact cells were washed twice with ice-cold DPBS and incubated with the membrane-impermeable biotinylation reagent Sulfo-NHS SS-Biotin (0.5 mg ⁄ mL; APExBIO, #A8005) in DPBS containing 0.1 mM CaCl_2_ and 1 mM MgCl_2_ (PBS+CM) for 30 min at 4 °C to label surface membrane proteins. To quench the reaction, cells were incubated with 10 mM glycine, pH 7.5 in ice-cold PBS+CM twice for 5 min at 4 °C. Sulfhydryl groups were blocked by incubating the cells with 5 nM N-ethylmaleimide (NEM) in DPBS for 15 min at room temperature. Cells were solubilized for 1 h at 4 °C in solubilization buffer (Triton X-100, 1%; Tris–HCl, 50 mM; NaCl, 150 mM; and EDTA, 5 mM; pH 7.5) supplemented with Roche complete protease inhibitor cocktail and 5 mM NEM. The lysates were cleared by centrifugation (20,000 × g, 10 min at 4 °C) to pellet cellular debris. The supernatant contained the biotinylated surface proteins. The concentration of the supernatant was measured using microBCA assay. Biotinylated surface proteins were affinity-purified from the above supernatant by incubating for 1 h at 4 °C with 100 μL of immobilized neutravidin-conjugated agarose bead slurry (Thermo Scientific, catalog #: 29200). The samples were then subjected to centrifugation (20,000 ×g, 10 min, at 4 °C). The beads were washed six times with solubilization buffer. Surface proteins were eluted from beads by boiling for 5 min with 200 μL of LSB ⁄ Urea buffer (2x Laemmli sample buffer (LSB) with 100 mM DTT and 6 M urea; pH 6.8) for SDS-PAGE and Western blot analysis.

### Automated patch-clamping with IonFlux Mercury 16 instrument

Whole-cell currents were recorded for HEK293T cells expressing GABA_A_ receptors. Automated patch clamping was performed on the Ionflux Mercury 16 instrument (Fluxion Biosciences, California), as previously described ^14^. Briefly, the extracellular solution (ECS) contained the following (in mM): 142 NaCl, 8 KCl, 6 MgCl_2_, 1 CaCl_2_, 10 glucose, and 10 HEPES; the intracellular solution (ICS) contained the following (in mM): 153 KCl, 1 MgCl_2_, 5 EGTA, and 10 HEPES. Cells were grown to 50 to 70 percent confluence on 10-cm dishes. On the day of experiments, cells were detached using accutase (Sigma Aldrich, catalog #: A6964-500mL) and suspended in serum free medium HEK293 SFM II (Gibco, catalog #: 11686-029), supplemented with 25 mM HEPES (Gibco, catalog #: 15630-080) and 1% penicillin streptomycin. Then cells were pelleted, resuspended in the ECS, and added to the Ionflux ensemble plate 16 (Fluxion Biosciences, catalog #: 910-0054). The ensemble plates were prepared according to manufacture suggestions. Whole-cell GABA-induced currents were recorded at a holding potential of −60 mV. The signals were acquired and analyzed by Fluxion Data Analyzer.

Dose-response curves elicited by GABA were fitted to the following equation using Origin software:

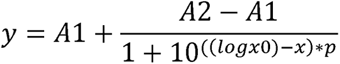

In the above equation, y is the normalized peak current, x is log(Concentration of GABA), x0 is the Half maximal effective concentration (EC_50_), and p is the Hill coefficient.

### Pixel-based sensitized acceptor emission FRET microscopy

Pixel-by-pixel based sensitized acceptor FRET microscopy was performed as described previously ^17, 64^. For FRET experiments on GABA_A_ receptors, (1) for FRET pair samples, HEK293T cells on coverslips were transfected with α1-CFP (donor) (0.5 μg), β2-YFP (acceptor) (0.5 μg), and γ2 (0.5 μg) subunits; (2) for the donor-only samples, to determine the spectral bleed-through (SBT) parameter for the donor, HEK293T cells were transfected with α1-CFP (donor) (0.5 μg), β2 (0.5 μg), and γ2 (0.5 μg) subunits; (3) for the acceptor-only samples, to determine the SBT parameter for the acceptor, HEK293T cells were transfected with α1 (0.5 μg), β2-YFP (acceptor) (0.5 μg), and γ2 (0.5 μg) subunits. Whole-cell patch-clamp recordings in HEK293T cells showed that fluorescently tagged GABA_A_ receptors have similar dose-response curves of GABA as compared to untagged ion channels ^17^. The coverslips were mounted using fluoromount-G (VWR, catalog #: 100502-406) and sealed. An Olympus Fluoview FV1000 confocal laser scanning system was used for imaging with a 60× 1.35 numerical aperture oil objective by using Olympus FV10-ASW software.

For the FRET pair samples, donor images were acquired at an excitation wavelength of 433 nm and an emission wavelength of 478 nm, FRET images at 433 nm excitation and 528 nm emission wavelengths, and acceptor images at 514 nm excitation and 528 nm emission wavelengths. For the donor-only samples, donor images were acquired at 433 nm excitation and 478 nm emission wavelengths, and FRET images at 433 nm excitation and 528 nm emission wavelengths. For the acceptor-only samples, FRET images were acquired at an excitation of 433 nm and an emission of 528 nm, and acceptor images at an excitation of 514 nm and an emission of 528 nm. Image analysis of FRET efficiencies was performed using the PixFRET plugin of the ImageJ software ^65^. The bleed-through was determined for the donor and the acceptor. With the background and bleed-through correction, the net FRET (nFRET) was calculated according to equation (1).

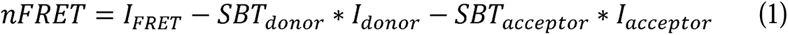

FRET efficiencies from sensitized emission experiments were calculated according to equation (2).

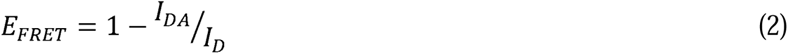

E_FRET_ represents FRET efficiency, I_DA_ represents the emission intensity of the donor in the presence of the acceptor, and I_D_ represents the emission intensity of the donor alone. Since I_D_ can be estimated by adding the nFRET signal amplitude to the amplitude of I_DA_ ^66^, FRET efficiency was calculated according to equation (3).

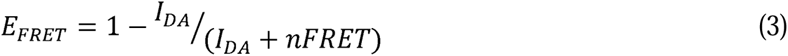

### Immunoprecipitation

Cell lysates (500 µg) were pre-cleared with 30 µl of protein A/G plus-agarose beads (Santa Cruz Biotechnology, catalog #: SC-2003) and 1.0 µg of normal mouse IgG (Santa Cruz Biotechnology, catalog #: SC-2025) for 1 hour at 4°C to remove nonspecific binding proteins. The pre-cleared cell lysates were incubated with 2.0 µg of mouse anti-α1 antibody for 1 hour at 4°C, and then with 30 µl of protein A/G plus agarose beads overnight at 4°C. The beads were then collected by centrifugation at 8000 ×g for 30 s, and washed three times with lysis buffer. The complex was eluted by incubation with 30 µl of Laemmli sample buffer in the presence of 5% 2-mercaptoethanol. The immunopurified eluents were separated in 8% SDS-PAGE gel, and Western blot analysis was performed.

### Quantitative RT-PCR

The relative gene expression levels were quantified using quantitative RT-PCR, as described previously ^15^. Briefly, cells were treated with chemicals for the indicate time. RNeasy Mini Kit (Qiagen, catalog #: 74104) was used to extract total RNA from cells. QuantiTect Reverse Transcription Kit (Qiagen, catalog #: 205311) was used to synthesize cDNA from total RNA. Quantitative PCR reactions (45 cycles of 15 s at 94°C (denaturation), 30 s at 59°C (annealing), and 30 s at 72°C (extension)) were performed using cDNA, PowerUp SYBR Green Master Mix (Applied Biosystems, catalog #: A25776) and corresponding primers in the QuantStudio 3 Real-Time PCR System (Applied Biosystems) and analyzed using QuantStudio software (Applied Biosystems). The forward and reverse primers for genes of interest and *RPLP2* (housekeeping gene control) are listed in **Supplementary Table S2**.

Threshold cycle (C_T_) was extracted from the PCR amplification plot, and the ΔC_T_ value was defined as:

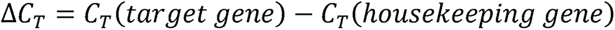

The relative mRNA expression level of target genes of drug-treated cells was normalized to that of untreated cells:

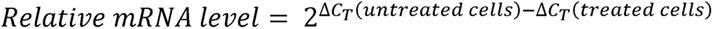

Each data point was evaluated in triplicate and measured three times.

### Cycloheximide-chase assay

HEK293T cells stably expressing α1(G251D)β2γ2 GABA_A_ receptors were seeded at 2.5 × 10^5^ cells per well in 6-well plates and incubated at 37°C overnight. Cells were then treated with Hispidulin or TP003 for 24 h prior to cycloheximide-chase assay. To stop protein translation, cells were treated with 100 μg/mL cycloheximide, a potent protein synthesis inhibitor (Enzo Life Sciences, catalog #: ALX-380-269). Cells were then chased for the indicated time, harvested, and lysed for SDS-PAGE and Western blot analysis.

### Generation of **α**1(G251D) knockin in human induced pluripotent stem cells (iPSCs)

The knockin of the G251D *GABRA1* variation (NM_001127644.2(GABRA1):c.752G>A (p.Gly251Asp); single nucleotide variant: rs1064793933) into apparently healthy male iPSCs (Applied StemCell, catalog #: ASE-9211) was generated using the CRISPR/Cas9 technique by Applied StemCell ^67^. To edit iPSCs, a ribonucleoprotein (RNP) was complexed using purified SpCas9 proteins (Integrated DNA Technologies) and two chemically synthesized guide RNA (gRNA) oligonucleotides (Integrated DNA Technologies) at optimized ratio in Nuclease-free Duplex Buffer (Integrated DNA Technologies, catalog#: 11-05-01-12). The sequences of two gRNAs are: AGCCAATCTTTCTCTTCAAG and TTCCACTTGAAGAGAAAGAT. The RNP was incubated at room temperature for 20 mins prior to electroporation with the 4D-Nucleofector System (Lonza). Next 1 × 10^6^ cells resuspended in 93 μL of P3 Primary Cell Nucleofector Solution (P3 Primary Cell 4D-Nucleofector X Kit, Lonza, Catalog #: V4XP-3024) were mixed with 7 μL of RNP and 12 μg of single-stranded oligodeoxynucleotide (ssODN) repair DNA template and then electroporated with the optimal pulse code. The sequence of ssODN is: CTTGTGTGTTTTACTTCTCAGGAGAATATGTTGTTATGACCACTCATTTCCATTTGAAGAGAAAAATTGACTACTTTGTTATTCAAACATACCTGCCATGCATAATGACAGTGATTCTCTCAC. Electroporated iPSCs were cultured 72-96 hrs before being sorted with Fluorescence Activated Cell Sorting (FACS) (BD Melody Cell Sorter) into multiple 96-well plates for subsequent clone screening via PCR amplification of gRNA target genomic regions and analyzed by Sanger Sequencing. The heterozygous knockin of the G251D *GABRA1* variation was confirmed by genotyping (**Supplementary Fig. S5**) using the following primers: forward, CAGATTTGTTCCTCCAATATGTGATG; reverse, CATATCTGTGACACTGCAAAGAG.

### GABAergic neuronal differentiation from human iPSCs

GABAergic neurons (iNs) were generated by expression of transcription factors ASCL1 and DXL2 using lentivirus transduction in iPSCs harboring the G251D *GABRA1* variation ^57^.

Lentivirus were generated from transiently transfected LentiX-HEK293T cells (Takara, catalog #: 632180) and collected after 60 hours from the media. Briefly, HEK293T cells were grown in 15-cm dishes and allowed to reach ~70-80% confluency before transient transfection using TransIT-2020 (Mirus), according to the manufacturer’s instruction. Each of the following plasmids (15 µg) was added to one 15-cm dish: ASCL1 (TetO-ASCL1-puro, Addgene plasmid # 97329), DXL2 (TetO-DXL2-hygro, Addgene plasmid # 97330), or rtTA expression vector (FUW-M2rtTA, Addgene plasmid # 20342). Additionally, to form the lentivirus, the following packaging and envelop plasmids were added to all of the 15-cm dishes as well: psPAX2 (15 µg) and pMD2.G (5 µg). TetO-Ascl1-puro (Addgene plasmid # 97329; http://n2t.net/addgene:97329; RRID: Addgene_97329) and DLX2-hygro (Addgene plasmid # 97330; http://n2t.net/addgene:97330; RRID: Addgene_97330) were a gift from Marius Wernig ^57^; FUW-M2rtTA was a gift from Rudolf Jaenisch (Addgene plasmid # 20342; http://n2t.net/addgene:20342; RRID:Addgene_20342) ^68^. The psPAX2 (Addgene plasmid # 12260) and pMD2.G (Addgene plasmid # 12259) were a gift from Didier Trono. Media was changed after 8 h; after 52 additional h, media was harvested and passed through 0.45 µm filter (Advantec, catalog #: 25CS045AS) to collect the lentivirus. Furthermore, the lentivirus were concentrated using Lenti-X concentrator (Takara Bio, catalog #: 631231) and quantified with the qPCR lentivirus titration kit (Abmgood, catalog #: LV900) according to the manufacturer’s instruction, and saved to −80°C for iPSCs transduction.

The 1 × 10^6^ iPSCs were plated onto a Matrigel (Corning, catalog #: 356277)-coated 6-cm dish in StemFlex media (Gibco, catalog #: A3349401) containing 10 μM Y-27632 (Biogems, catalog #1293823) with an assembly of lentivirus (ASCL1, DXL2, and rtTA lentivirus). Twenty-four hours post infection, the virus-containing media was removed and N2 media with doxycycline (DOX, 2 µg/mL, Sigma, catalog #: D9891) was added. N2 media contains 1% of N2 supplement (ThermoFisher, catalog #: 17502048) in DMEM/F12 (ThermoFisher, catalog #: 11320-082). N2 media containing DOX was changed daily; proper selection reagents were added 48 hours post infection: ASCL1 expressing cells were selected with 1 µg/mL puromycin (DOT scientific, catalog #: DSP33020) and DLX2 expressing cells with 100 µg/mL hygromycin (ThermoFisher, catalog #: J60681-03). The infection efficiency was measured 72 hours post infection, ie., 48 h after the addition of DOX, using immunofluorescence with appropriate antibodies for each factor. Optimal reprogramming efficiencies were achieved with greater than 80% of cells expressing ASCL1 and DLX2. Upon obtaining optimal reprogramming efficiencies, subsequently similarly, infected iPSCs were allowed to differentiate in N2 media, with appropriate DOX, puromycin, and hygromycin. Cytosine arabinose (Ara C, Sigma, catalog #: C6645) was added at a final concentration of 4 µM from DPI 5 (day-post-infection).

On DPI 7, iN cells (1 × 10^4^ per well) were plated together with mouse glial cells (2 × 10^4^ per well) onto 8-well chamber glass slides (ThermoFisher, catalog #: 154941) in the Growth Medium. Prior to plating cells, the slides were coated with 10 μg/mL poly-D-lysine (PDL, Sigma, catalog #: P6407) at 4°C overnight. The next day PDL was removed and the slides were washed with sterile distilled water three times. Then they were coated with 10 µg/mL Laminin (Sigma, catalog #: L2020) in a 37°C cell incubator for at least 2 hours. Laminin was removed immediately before plating cells. The Growth Medium contained the following: 100 mL Neurobasal A (Gibco, catalog #: 21103049), 2 mL B27 (Gibco, catalog #: 17504044), 1 mL GlutaMAX (Gibco, catalog #: 35050061), 2.5 mL of heat-inactivated fetal bovine serum (ThermoFisher, catalog #: SH3039603HI) and 1 mL Penicillin-Streptomycin (Hyclone, catalog #: sv30010). The Growth Medium contained DOX until DPI 14; from DPI 15 onward, the following were also supplemented: 20 ng/mL brain-derived neurotrophic factor (BDNF; R&D systems, catalog #: 248-BD), 20 ng/mL glial cell line-derived neurotrophic factor (GDNF; R&D systems, catalog #: 212-GD), and 10 µM ROCK inhibitor Y27632 (HelloBio, catalog #: HB2297). Additionally, when the glial cells grow to ~80-90% confluent, Ara C was added at a final concentration of 4 µM. The medium was replaced every 3-4 days. After 3-4 weeks post infection, GABAergic neurons were subjected to immunofluorescence staining or *In situ* Proximity Ligation Assay (PLA) for confocal microscopy.

### Immunofluorescence Staining and Confocal Microscopy

Neuron staining and confocal immunofluorescence microscopy analysis were performed as described previously ^14^. Briefly, to label cell surface proteins, iPSC-derived GABAergic neurons on the chamber slides were fixed with 4% paraformaldehyde in DPBS for 10 min. We then blocked with 10% goat serum (ThermoFisher, catalog #: 16210064) in DPBS for 0.5 h, and without detergent permeabilization, incubated with 100 μL of appropriate primary antibodies against the GABA_A_ receptor α1 subunit (Synaptic Systems, Goettingen, Germany, catalog #: 224203) (1:250 dilution), β2/3 subunit (Millipore, catalog #: 05-474) (1:250 dilution), or γ2 subunit (Synaptic Systems, Goettingen, Germany, catalog #: 224003) (1:250) diluted in 2% goat serum in DPBS, at room temperature for 1 h. Then the neurons were incubated with Alexa 594-conjugated goat anti-rabbit antibody (ThermoFisher, catalog #: A11037), or Alexa 594-conjugated goat anti-mouse antibody (ThermoFisher, catalog #: A11032) (1:500 dilution) diluted in 2% goat serum in DPBS for 1 h. Afterward, the chamber slides were mounted using fluoromount-G (VWR, catalog #: 100502-406) and sealed. An Olympus IX-81 Fluoview FV3000 confocal laser scanning system was used. A 60× 1.40 numerical aperture oil objective was used to collect high-resolution images using FV31S-SW software. The images were analyzed using ImageJ software ^69^.

### *In situ* Proximity Ligation Assay (PLA) and quantification of PLA signal

GABAergic neurons (iNs) grown on the chamber slides were fixed with 4% paraformaldehyde (PFA, ThermoFisher, catalog #: 28908) for 15 min at room temperature, followed by quenching twice with 100 mM glycine in DPBS for 5 min. Then iNs were washed gently twice with DPBS and permeabilized with 0.2% saponin (VWR, catalog #: 80058-63) for 15 min. Afterwards, iNs were subjected to *in situ* PLA using Duolink In Situ Detection Reagents Red kit (Sigma, catalog #: DUO92008) according to manufacturer’s instructions. As indicated, incubations at 37°C were performed in a humidity chamber or sealed with Parafilm. Briefly, first, iNs were blocked with the blocking reagent at 37°C for 1 hour, and then incubated with two primary antibodies against GABA_A_ receptor α1 subunit and another interacting protein; these antibodies are raised in different species (mouse and rabbit) and their epitopes are in the same cellular compartment of ER lumen. Primary antibodies were diluted in the antibody diluent (DUO82008) at a ratio of 1:250 and incubated overnight at 4°C. Secondly, slides were washed gently twice with DPBS and then incubated with the PLA Probes of Anti-mouse MINUS (DUO92004) and Anti-rabbit PLUS (DUO92002) for 1 hour at 37°C. Thirdly, slides were washed gently twice with 1× Buffer A (DUO82049) and incubated with the DNA ligase in the Ligation buffer for 30 min at 37°C. Fourthly, slides were washed gently twice with 1× Buffer A and incubated with the DNA polymerase in the Amplification buffer for 100 min at 37°C. Finally, slides were washed gently twice with 1× Buffer B, followed by once with 0.01× Buffer B, and then mounted with mounting medium with DAPI (DUO82040) to stain the nuclei. The detection probes contain a red fluorophore of 594 nm excitation.

For confocal microscopy, an Olympus IX-81 Fluoview FV3000 confocal laser scanning system was used. A 60× 1.40 numerical aperture oil objective was used to collect high-resolution images using FV31S-SW software. Quantification of the PLA puncta was achieved using the ImageJ software from the NIH with the built-in macro *Analyze Particles* ^70^. Images were smoothed; a threshold was selected manually to account for PLA puncta from background fluorescence, and such threshold was applied to all images in the sample set. Objects larger than 5 μm^2^ were excluded such as nuclei, and the built-in macro *Analyze Particles* was used to count PLA puncta per cell.

### Statistical analysis

All data are presented as mean ± SEM. Statistical significance was evaluated using a two-tailed Student’s t-test between two groups as appropriate, and the analysis of variance (ANOVA) followed by a post-hoc Tukey test was applied for comparisons in multiple groups. A *p* < 0.05 was considered statistically significant. ∗, p< 0.05; ∗∗, p< 0.01; ∗∗∗, p< 0.001.

## Acknowledgements

This work was supported by the National Institutes of Health (R01NS105789 and R01NS117176 to TM, R01NS123524 to AS, and T32GM135081 to LA) and the Brain Research Foundation (BRFSG-2021-08 to AS).

## Author contributions

Conceptualization, YW and TM; Data curation: YW, HS, and LA; Formal analysis: YW, HS, and TM; Funding acquisition: AS, LA, and TM; Supervision: TM; Writing – original draft: YW and TM; Writing – review & editing: YW, HS, LA, AS, and TM.

## Supplementary information

Supplementary information includes five supplementary figures and two supplementary tables.

**Fig S1**. **Cartoon representation of GABA_A_ receptors. a** Pentameric α1β2γ2 receptors were constructed from their cryo-EM structure (6X3S.pdb) with side view (left) and the top view from the synaptic cleft. **b** Binding sites of GABA and benzodiazepines in the subunit-subunit interfaces.

**Fig S2**. ***In silico* saturation mutagenesis analysis of the human** α**1 subunit of GABA_A_ receptors.** Rhapsody (http://rhapsody.csb.pitt.edu) was used to predict the pathogenic probability of all possible 19 amino acid substitutions at each site on the sequence of human α1 subunit of GABA_A_ receptors (P14867), based on the protein’s sequence, structure, and dynamics. The signature cys-loop, loops A-F that form the ligand-binding pockets, α-helices (α1, α2) and β sheets (β1-β10) in the NTD, and transmembrane helices (M1-M4) were labelled. Known sites that carry missense clinical variants were colored in red. The sites of eight selected variants in this study were highlighted in green.

**Fig S3**. **Screening of known GABA_A_ receptor modulators reveals pharmacological chaperones that increase the total protein level of** α**1 DAVs.** HEK293T cells stably expressing epilepsy-associated α1(D219N)β2γ2 (**a**), α1(G251D)β2γ2 (**b**), or α1(P260L)β2γ2 (**c**) GABA_A_ receptors were treated with indicated chemicals (10 μM) for 24 h. Then cells were lysed, and total proteins were subjected to SDS-PAGE and Western blot analysis. BiP was detected as an indication of the ER stress. β-actin served as a protein loading control. Quantification of the normalized α1 band intensity was shown on bottom panels (*n* = 3). Each data point is reported as mean ± SEM. One-way ANOVA followed by post-hoc Tukey test was used for statistical analysis. * *p* < 0.05; ** *p* < 0.01; *** *p* < 0.001.

**Fig S4**. **Effect of Hispidulin and TP003 on the total protein level of the wild type** α**1 subunit.** HEK293T cells stably expressing wild type α1β2γ2 GABA_A_ receptors were treated with Hispidulin (0.05 μM to 25 μM, 24 h) (**a**) or TP003 (0.05 μM to 10 μM, 24 h) (**b**). Then cells were lysed, and total proteins were subjected to SDS-PAGE and Western blot analysis. BiP was detected as an indication of the ER stress. β-actin served as a protein loading control. Quantification of the normalized α1 band intensity was shown on bottom panels (*n* = 3). Each data point is reported as mean ± SEM. One-way ANOVA followed by post-hoc Tukey test was used for statistical analysis. * *p* < 0.05.

**Fig S5. Sanger sequencing to confirm the heterozygous knockin of α1(G251D) in human iPSCs.**

**Table S1.** Predicted pathogenic probability of known clinical variants of the human α1 subunit of GABA_A_ receptors according to Rhapsody.

**Table S2.** List of qPCR primers.

