## supplemental Figures for "Pharmacological chaperones restore proteostasis of epilepsy-associated GABA_A_ receptor variants"

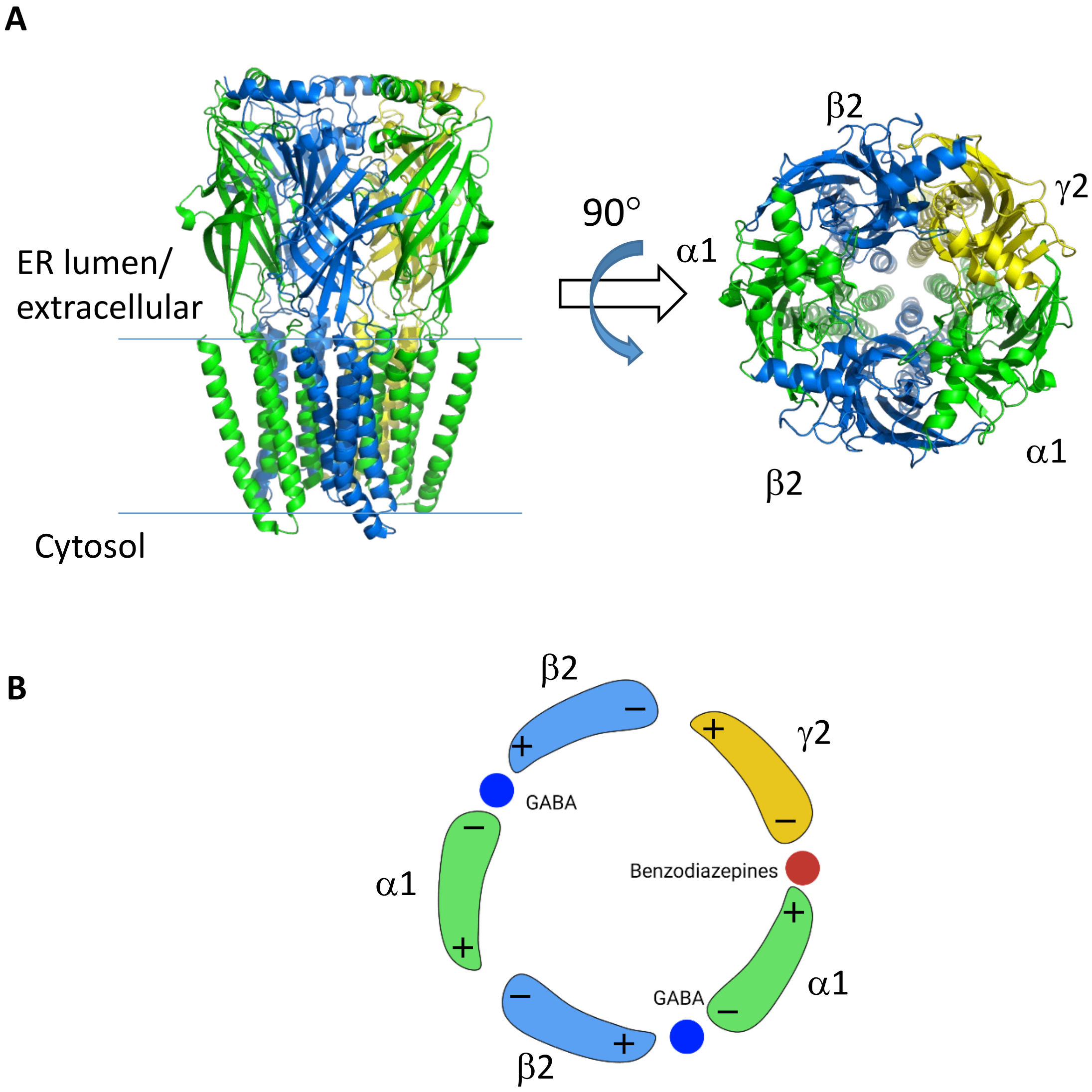


**Supplementary Figure S1**. **Cartoon representation of GABA$_A$ receptors. a** Pentameric α1β2γ2 receptors were constructed from their cryo-EM structure (6X3S.pdb) with the side view (left) and the top view from the synaptic cleft. **b** Binding sites of GABA and benzodiazepines in the subunit-subunit interfaces.

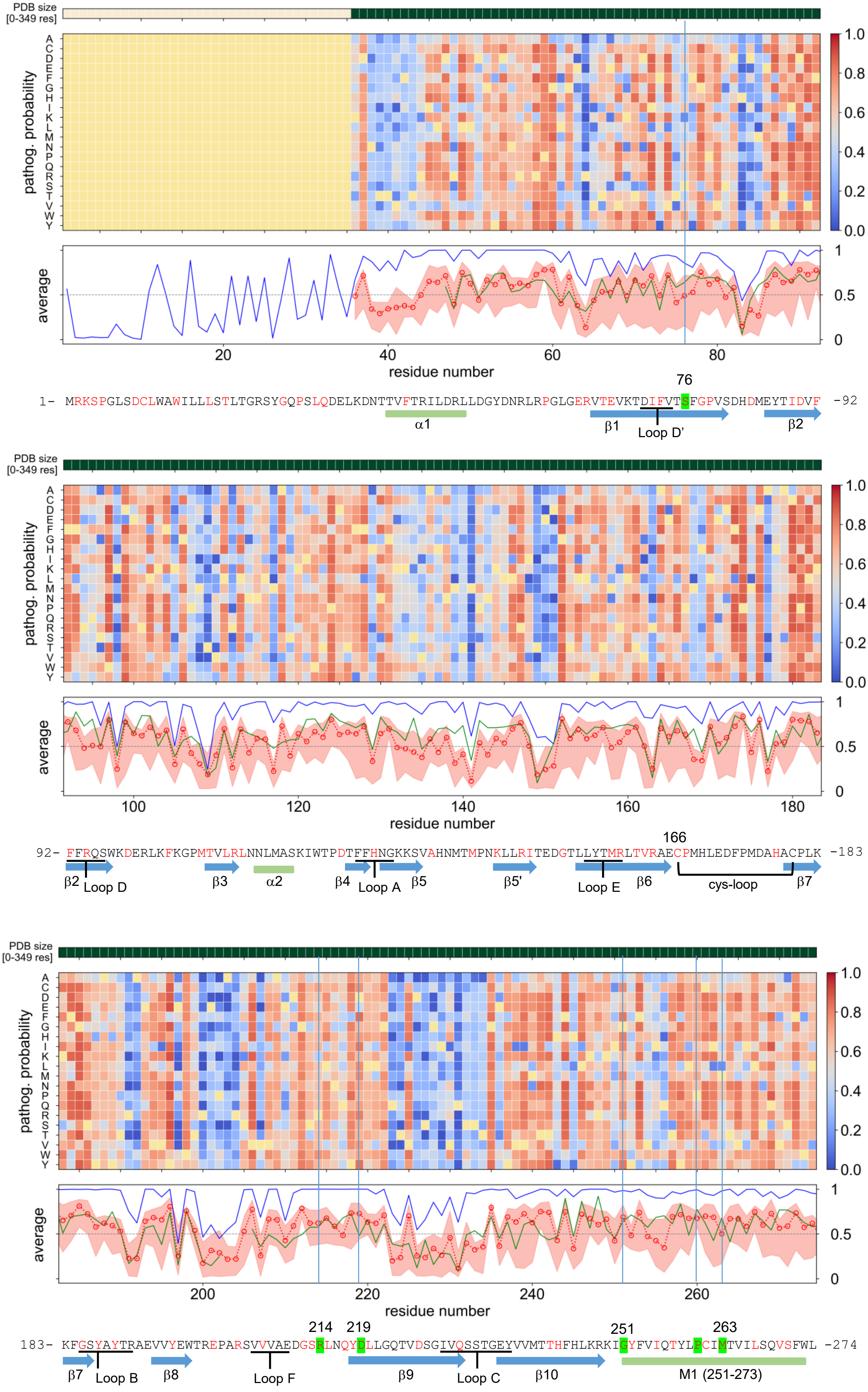

PDB size [0-349 res]
pathog. probability
A C D E F G H I K L M N P Q R S T V W Y
1.0 0.8 0.6 0.4 0.2 0.0
average
1 0.5 0
20 40 60 80
residue number
76
1- MRKSPGLSDCLWAWILLLSTLTGRSYGQPSLQDELKDNTTVFTRILDRLLDGYDNRLRPGLGERVTEVKTDIFVTSFGPVSDHDMEYTIDVF -92
α1
β1
Loop D'
β2
PDB size [0-349 res]
pathog. probability
average
100 120 140 160 180
residue number
166
92- FFRQSWKDERLKFKGPMTVLRLNNLMASKIWTPDTFFHNGKKSVAHNMTMPNKLLRITEDGTLLYTMRLTVRAECPMHLEDFPMDAHACPLK -183
β2
Loop D
β3
α2
β4
Loop A
β5
β5'
Loop E
β6
cys-loop
β7
PDB size [0-349 res]
pathog. probability
average
200 220 240 260
residue number
214
219
251
263
183- KFGSYAYTRAEVVYEWTREPARSVVVAEDGSRLNQYDLLGQTVDSGIVQSSTGEYVVMTTHFHLKRKIGYFVIQTYLPCIMTVILSQVSFWL -274
β7
Loop B
β8
Loop F
β9
Loop C
β10
M1 (251-273)

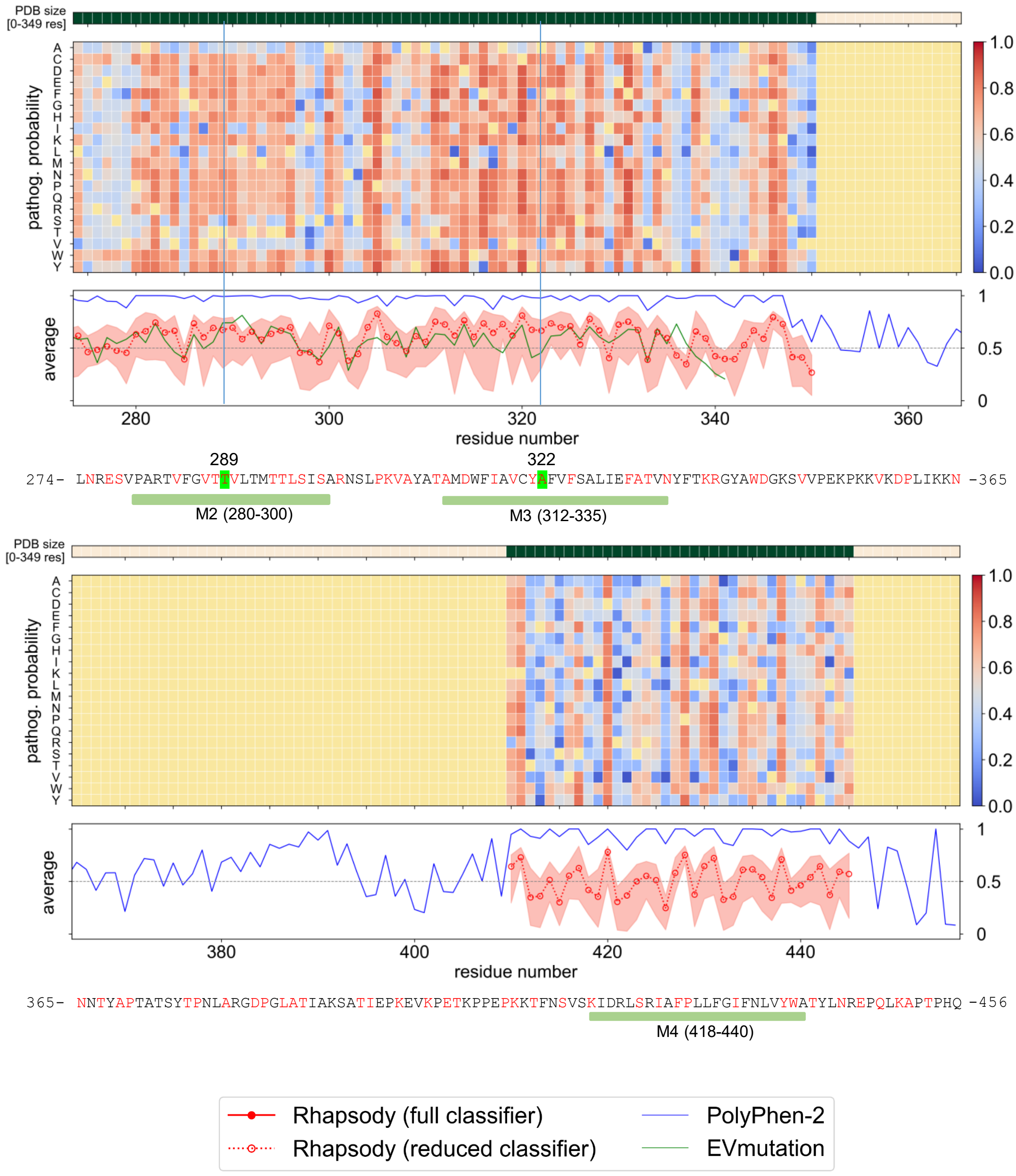


**Supplementary Figure S2. *In silico* saturation mutagenesis analysis of the human α1 subunit of GABA$_A$ receptors.** Rhapsody (http://rhapsody.csb.pitt.edu) was used to predict the pathogenic probability of all possible 19 amino acid substitutions at each site on the sequence of human α1 subunit of GABA$_A$ receptors (P14867), based on the protein's sequence, structure, and dynamics. The signature cys-loop, loops A-F that form the ligand-binding pockets, α-helices (α1, α2) and β sheets (β1-β10) in the NTD, and transmembrane helices (M1-M4) were labelled. Known sites that carry missense clinical variants were colored in red. The sites of eight selected variants in this study were highlighted in green.

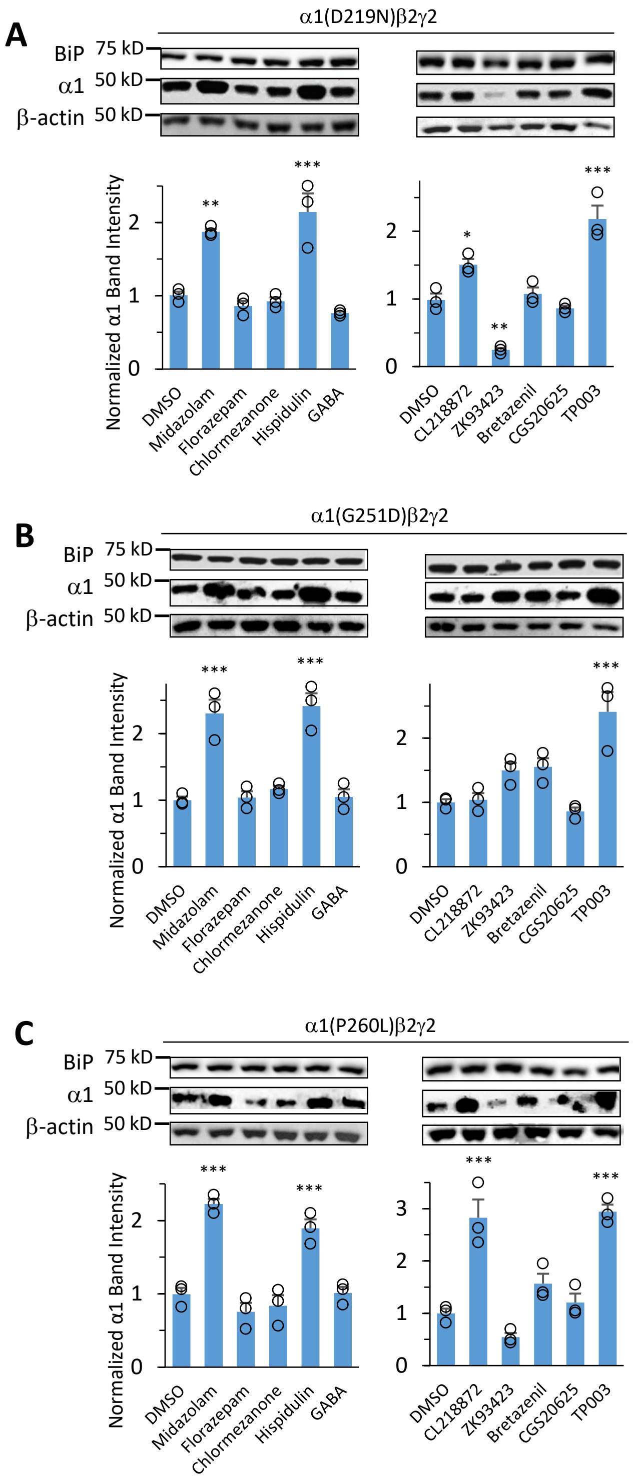


**Supplementary Figure S3. Screening of known $GABA_A$ receptor modulators reveals pharmacological chaperones that increase the total protein level of α1 DAVs.** HEK293T cells stably expressing epilepsy-associated α1(D219N)β2γ2 (**a**), α1(G251D)β2γ2 (**b**), or α1(P260L)β2γ2 (**c**) $GABA_A$ receptors were treated with indicated chemicals (10 μM) for 24 h. Then cells were lysed, and total proteins were subjected to SDS-PAGE and Western blot analysis. BiP was detected as an indication of the ER stress. β-actin served as a protein loading control. Quantification of the normalized α1 band intensity was shown on bottom panels ($n$ = 3). Each data point is reported as mean ± SEM. One-way ANOVA followed by post-hoc Tukey test was used for statistical analysis. * $p < 0.05$; ** $p < 0.01$; *** $p < 0.001$.

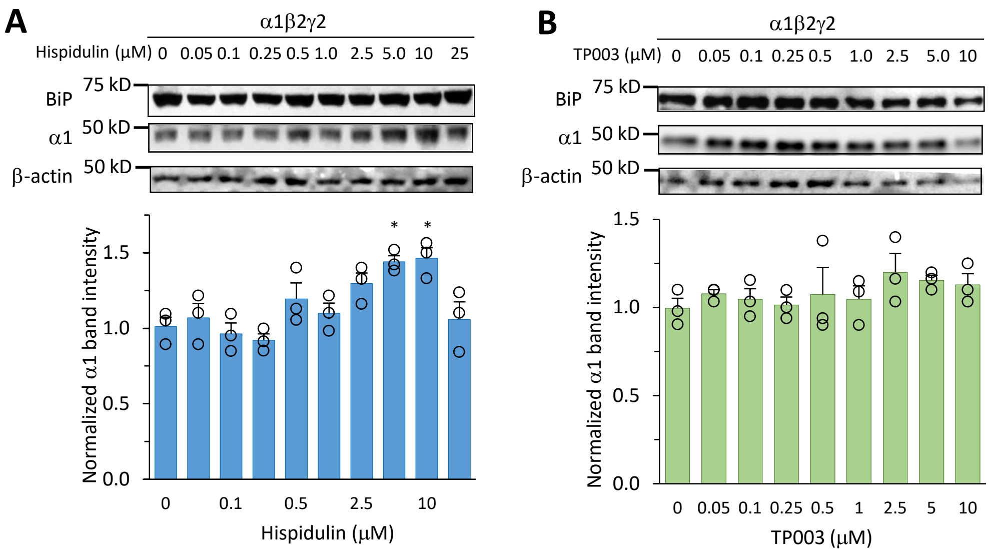


**Supplementary Figure S4. Effect of Hispidulin and TP003 on the total protein level of the wild type α1 subunit.** HEK293T cells stably expressing wild type α1β2γ2 $GABA_A$ receptors were treated with Hispidulin (0.05 μM to 25 μM, 24 h) (**a**) or TP003 (0.05 μM to 10 μM, 24 h) (**b**). Then cells were lysed, and total proteins were subjected to SDS-PAGE and Western blot analysis. BiP was detected as an indication of the ER stress. β-actin served as a protein loading control. Quantification of the normalized α1 band intensity was shown on bottom panels ($n = 3$). Each data point is reported as mean ± SEM. One-way ANOVA followed by post-hoc Tukey test was used for statistical analysis. * $p < 0.05$.

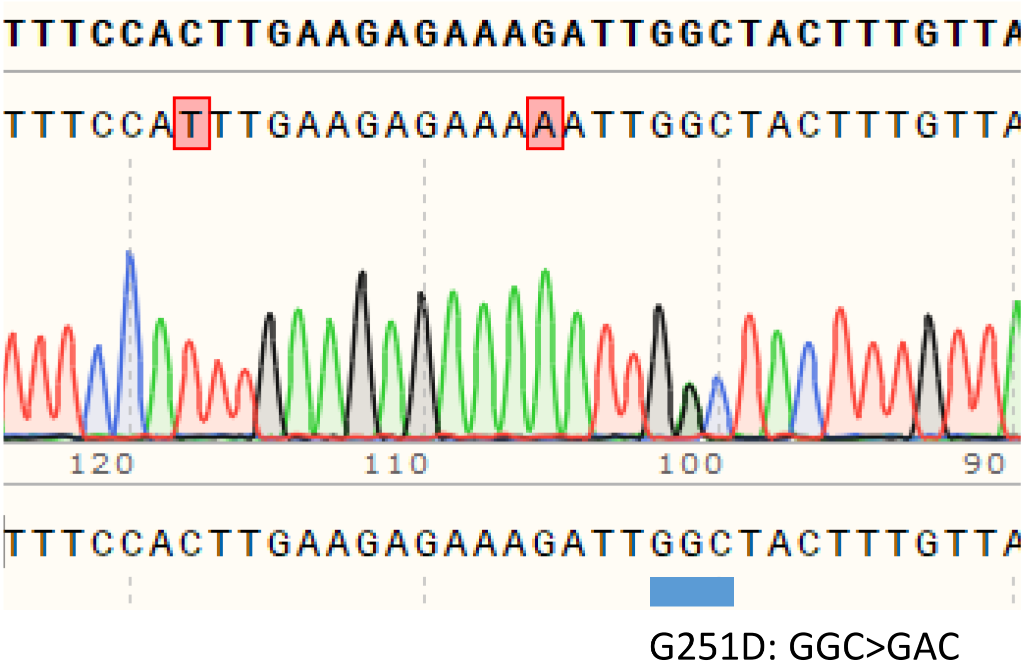


**Supplementary Figure S5**. **Sanger sequencing to confirm the heterozygous knockin of α1(G251D) in human iPSCs.**
